# Queuine is a nutritional regulator of *Entamoeba histolytica* response to oxidative stress and a virulence attenuator

**DOI:** 10.1101/2020.04.30.070276

**Authors:** Shruti Nagaraja, Maggi W. Cai, Jingjing Sun, Hugo Varet, Lotem Sarid, Meirav Trebicz-Geffen, Yana Shaulov, Mohit Mazumdar, Rachel Legendre, Jean-Yves Coppée, Thomas J. Begley, Peter C. Dedon, Samudrala Gourinath, Nancy Guillen, Yumiko Saito-Nakano, Chikako Shimokawa, Hajime Hisaeda, Serge Ankri

## Abstract

Queuosine is a naturally occurring modified ribonucleoside found in the first position of the anticodon of the transfer RNAs for Asp, Asn, His and Tyr. Eukaryotes lack pathways to synthesize queuine, the nucleobase precursor to queuosine, and must obtain it from diet or gut microbiota. Here we describe the effects of queuine on the physiology of the eukaryotic parasite, *Entamoeba histolytica*, the causative agent of amebic dysentery. Queuine is efficiently incorporated into *E. histolytica* tRNAs by a tRNA-guanine transglycosylase (EhTGT) and this incorporation stimulates the methylation of C_38_ in tRNA^Asp^_GUC_. Queuine protects the parasite against oxidative stress (OS) and antagonizes the negative effect that oxidation has on translation by inducing the expression of genes involved in OS response, such as heat shock protein 70 (Hsp 70), antioxidant enzymes, and enzymes involved in DNA repair. On the other hand, queuine impairs *E. histolytica* virulence by downregulating the expression of genes previously associated with virulence, including cysteine proteases, cytoskeletal proteins, and small GTPases. Silencing of EhTGT prevents incorporation of queuine into tRNAs and strongly impairs methylation of C_38_ in tRNA^Asp^_GUC_, parasite growth, resistance to OS, and cytopathic activity. Overall, our data reveal that queuine plays a dual role in promoting OS resistance and reducing parasite virulence.

**Importance:** *Entamoeba histolytica* is a unicellular parasite that causes amebiasis. The parasite resides in the colon and feeds on the colonic microbiota. The gut flora is implicated in the onset of symptomatic amebiasis due to alterations in the composition of the bacteria. These bacteria modulate the physiology of the parasite and affect the virulence of the parasite through unknown mechanisms. Queuine, a modified nucleobase of queuosine, is exclusively produced by the gut bacteria and leads to tRNA modification at the anticodon loops of specific tRNAs. We found that queuine induces a mild oxidative stress resistance in the parasite and attenuates its virulence. Our study highlights the importance of bacterially derived products in shaping the physiology of the parasite. The fact that queuine inhibits the virulence of *E. histolytica* may lead to new strategies for preventing and/or treating amebiasis by providing to the host queuine directly or via probiotics.

## Introduction

Amebiasis is an enormous global medical problem due to poor sanitary conditions and unsafe hygiene practices in many parts of the world. According to the World Health Organization, 50 million people in India, Southeast Asia, Africa, and Latin America suffer from amebic dysentery, with amebiasis causing the death of at least 100,000 people each year. *Entamoeba histolytica*, the etiologic agent of amebiasis, proliferate in the intestinal lumen and phagocytose resident gut flora. Over the last few decades, it has become evident that *E. histolytica’s* pathogenicity is directly linked to the parasite’s interaction with the gut microbiota [1]. This interaction is very selective because only those bacteria with the appropriate recognition molecules are ingested by the parasite [2]. It has been reported that the *E. histolytica*’s association with specific intestinal bacteria changes the parasite’s cell surface architecture [3, 4]. It has also been reported that phagocytosis of pathogenic bacteria boosts *E. histolytica*’s cytopathogenicity, increases the expression of Gal/GalNAc lectin on the cell surface, and boosts cysteine proteinase activity when trophozoites are co-cultured with the enteropathogenic *Escherichia coli* (EPEC) O55 or *Shigella dysenteriae* [5]. It has also been reported that bacteria-induced augmentation of *E. histolytica*’s virulence is achieved only when the trophozoites phagocytose intact live cells [6]. The composition of the gut flora in patients suffering from amebiasis shows a significant decrease in the population size of *Bacteroides, Clostridium coccoides, Clostridium leptum, Lactobacillus*, and *Campylobacter* and an increase in *Bifidobacterium*, while there was no change in *Ruminococcus* compared to healthy patients [7]. These findings suggest that the pathogenesis of amebiasis might be driven by a dysregulated microbiome or crosstalk between enteropathic bacteria, the parasite, and the intestinal immune system. This crosstalk may be modulated by chemical signaling molecules, such as short-chain fatty acids (SCFAs) released by the bacteria [8], or by bacterial oxaloacetate that regulates parasite virulence and resistance to oxidative stress (OS) [9, 10]. The modified ribonucleoside, queuosine, 7-(((4.5-cis-dihydroxy-2-cyclopenten-1-yl)-amino)-methyl)-7-deazaguanosine) may also participate in the crosstalk between the parasite and the microbiota. Eubacteria can synthesize the queuine nucleobase precursor of queuosine *de novo*. However, a recent study showed that some Eubacteria can also salvage precursors of queuosine [11]. Since mammals cannot synthesize queuine and salvage it from diet and gut microbes, these observations suggest a dynamic supply and demand for queuine by both gut microbiota and the host [12–14]. The diverse roles of queuosine in bacterial physiology are only now emerging. tRNA-guanine transglycosylase (TGT), which catalyzes the exchange of queuine for guanine at the wobble position of certain tRNAs, has been recognized as one of the key enzymes that regulate virulence in *Shigella flexneri* [15, 16]. Recently, it has been found that a modification of the wobble position of tRNA with a GUN anticodon by 7-deaza-guanosine derivative queuosine (Q34) stimulates *Schizosaccharomyces pombe* Dnmt2/Pmt1-dependent C38 methylation (m^5^C_38_) in the tRNA^Asp^_GUC_ anticodon loop [17], with the latter correlated with resistance to OS in many organisms including *E. histolytica* [18, 19]. The fact that both methylation of C_38_-tRNA^Asp^_GUC_ and queuinosylation of tRNAs occur at the same anticodon loop suggests coordination among these modifications in regulating OS resistance in the parasite. It has been previously reported that queuine promotes the resistance of cancer cells to OS by increasing the activity of antioxidant enzymes [20]. In contrast, queuosine deficiency in tRNAs leads to the accumulation of misfolded proteins that induce a cellular stress response [17]. It has also been found that queuine enhances translational speed and fidelity in eukaryotes [21].

Here we report that queuine: (i) induces C_38_ hypermethylation (m^5^C_38_) in the tRNA^Asp^_GUC_ anticodon loop, (ii) promotes resistance of the parasite to OS by triggering the expression of genes associated with stress response, (iii) restores protein synthesis in parasite exposed to OS, and (iv) reduces parasite virulence by reducing expression of genes associated with virulence.

## Results

### Queuine is incorporated into *E. histolytica* tRNA and induces C38 hypermethylation (m^5^C_38_) in the tRNA^Asp^_GUC_ anticodon loop

Given the established role of queuine as a precursor to Q in tRNA, we first set out to characterize the incorporation of queuine into amoebic tRNA. Here we quantified Q in tRNA in the parasite using LC-MS/MS. tRNA samples were prepared from untreated trophozoites and from trophozoites that were grown with queuine (0.1μM). This concentration of queuine has been chosen because (i) it provides to the parasite the best protection against H_2_O_2_ (Fig S1) and (ii) because it has also been used in a study on the social amoeba *Dictostelium discoideum* [22]. The amount of queuine in the human large intestine is unknown. However, its amount in the blood can reach 10 nM [23]. A 10 to 1000-time level of difference can be normally observed between the level of a metabolite in the large intestine versus its level in the blood [24]. Therefore, the concentration of queuine used in this study (0.1μM) is probably biologically relevant.

We observed that supplementation of queuine in the Trypticase Yeast Extract Iron Serum (TYI-S-33) growth medium caused a more than 5-fold increase in the level of Q in tRNAs as measured by LC-MS/MS (Fig 1A). We have also quantified the level of Q into tRNA^His^_GUG_ and tRNA^Asp^_GUC by_ using N-acryloyl-3-aminophenylboronic acid (APB) polyacrylamide gel or acidic polyacrylamide gel respectively to separate Q-tRNAs from unmodified tRNAs (Fig 2A & 2C). We observed a very strong increase of the level of Q-tRNA^His^_GUG_ and Q-tRNA^Asp^_GUC_ in trophozoites that were grown with queuine when compared to control trophozoites that were grown without queuine (Fig 2B & 2D).

**Figure 1:**
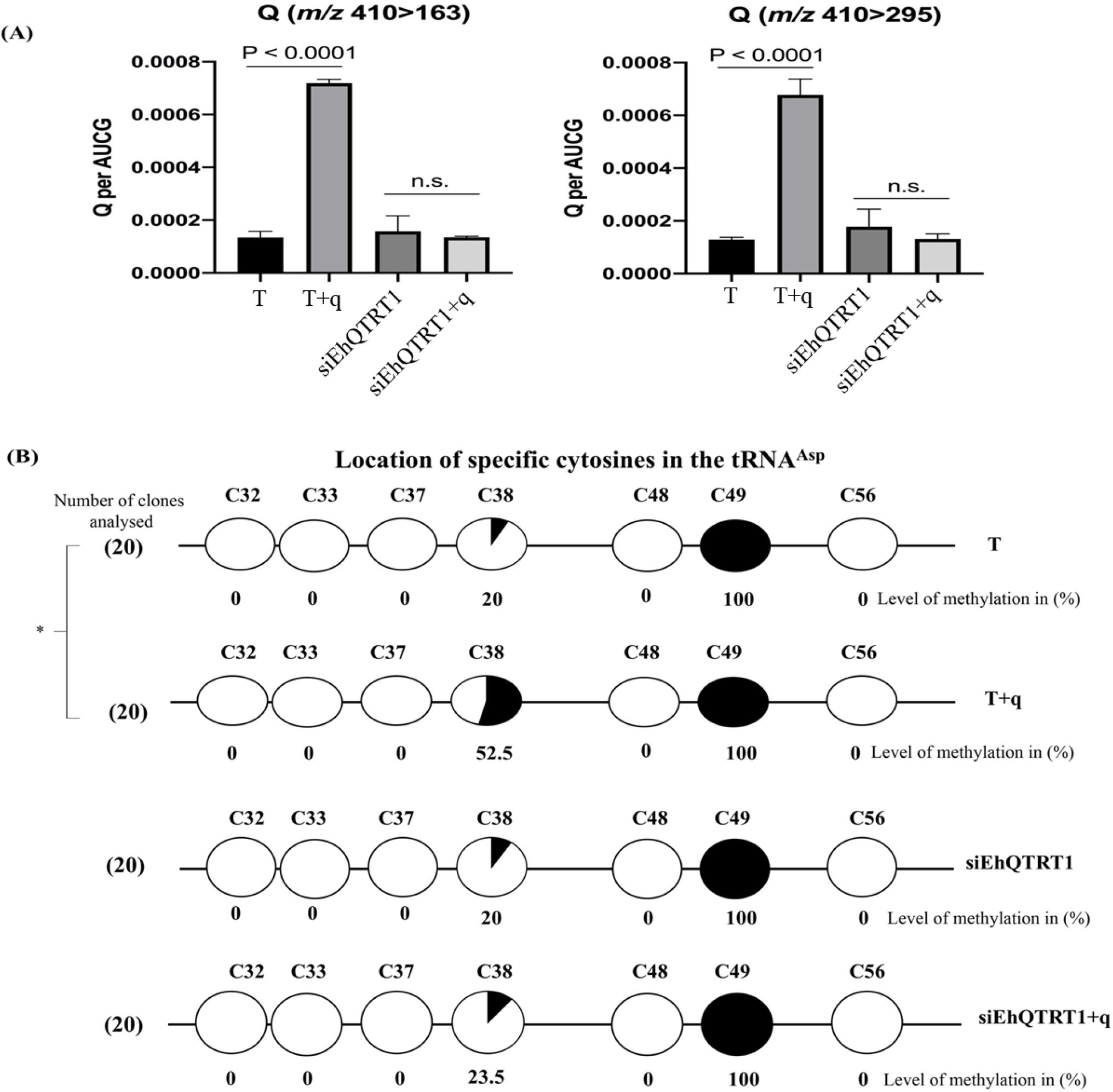
Queuine is efficiently incorporated into *E. histolytica* tRNA and leads to C38 hypermethylation (m^5^C_38_) in the anticodon loop of tRNA^Asp^_GUC_. (A) Quantification of Q in tRNA of *E. histolytica* trophozoites using LC-MS/MS. RNA samples were prepared from control trophozoites (T), trophozoites that were grown with queuine (q) supplemented in the culture medium (0.1 μM for 3 days) (T+q), trophozoites silenced for the expression of EhQTRT1 (siEhQTRT1) and trophozoites silenced for the expression of EhQTRT1 that were grown with queuine (siEhQTRT1+q). tRNA was purified and analyzed by LC-MS/MS to quantify Q using two transitions: *m/z* 410 → 163 (left) and *m/z* 410 → 295 (right). Data represent mean ± SEM for N=12 (Unpaired Student’s t-test, p≤0.0001). B) Bisulfite sequencing of tRNA^Asp^ GUC has been performed on control trophozoites (T), trophozoites that were grown with 0.1μM queuine for three days (T+q), trophozoites silenced for the expression of EhQTRT1 (siEhQTRT1) and trophozoites silenced for the expression of EhQTRT1 that were grown with queuine (siEhQTRT1+q). The number of independent clones (sequence reads) have been shown in parentheses towards the left side of each row. The black circles represent methylated cytosine residues, whereas the white circles represent unmethylated cytosine residues. The percentage of cytosine methylation is indicated below each circle. The location of specific cytosine residues has been indicated above the circles. The level of cytosine-38 tRNA^Asp^_GUC_ methylation in T+q (52.5± 3.5%;) was significantly different from that of T (20 ± 1.4%) according to the analysis with an unpaired Student’s t test (p<0.05). Indeed, the level of cytosine-38 tRNA^Asp^_GUC_ methylation in siEhQTRT1 trophozoites (siEhQTRT1) and in siEhQTRT1 trophozoites that were grown with queuine (siEhQTRT1+q) was not significantly different according to the analysis with an Unpaired Student’s t test (p<0.05). The data are expressed are mean ± SD obtained from 20 clones each.

**Figure 2:**
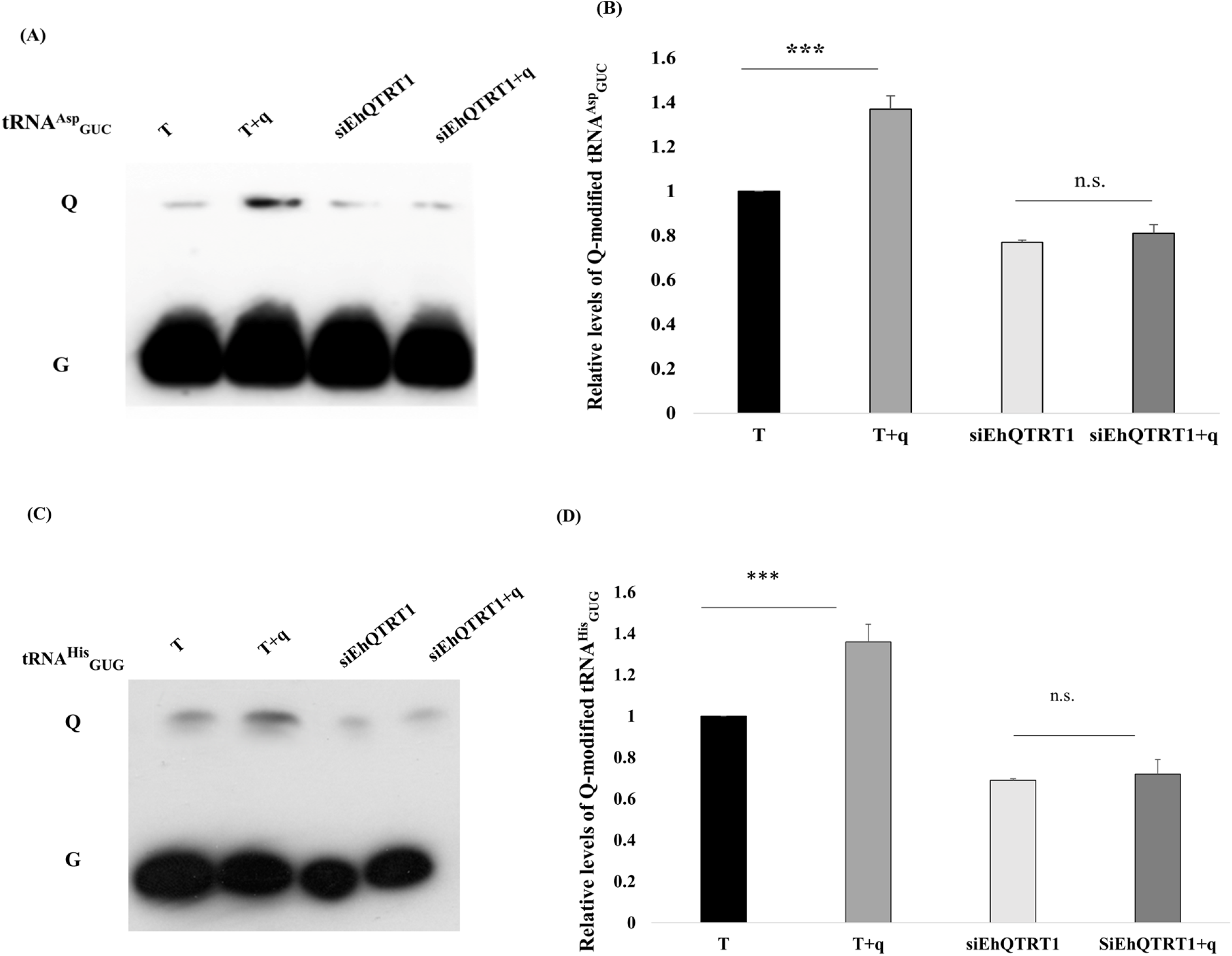
Acidic Urea-PAGE analysis of tRNA^Asp^_GUC_ and APB northern blot analysis of tRNA^His^_GUG_. (A) Acidic Urea-PAGE analysis of tRNA^Asp^_GTC_ in *E. histolytica* trophozoites. (A). Lane 1-Control trophozoites (T), Lane 2-Control trophozoites that were grown with queuine (T+q), Lane 3-siEhQTRT1 trophozoites (siEhQTRT1), and Lane 4-siEhQTRT1 trophozoites that were grown with queuine (siEhQTRT1+q). Total RNA samples was run on a 15% acrylamide gel supplemented with 0.1M sodium acetate pH 4.8, transferred on a nylon membrane, dried and incubated with a biotinylated probe against tRNA^Asp^_GUC_. (B) Quantitative analysis of relative levels of Q-modified tRNA^Asp^_GUC_. The signal intensities of the Q-tRNA band for each sample was divided by the signal intensities for their total tRNA content (Q+G) and then normalized to the wild type (T) samples. The data represent mean ± SD for three independent experiments (Unpaired Student’s t test, p = 0.0004) (C) APB northern blot analysis for tRNA^His^_GTG_. Lane 1-Control Trophozoites (T), Lane 2-Control trophozoites that were grown with queuine (T+q), Lane 3-siEhQTRT1 trophozoites that were grown with queuine (siEhQTRT1+q), and Lane 4-siEhQTRT1 trophozoites. Total RNA samples were run on a 15% acrylamide gel supplemented with APB, transferred on a nylon membrane, dried and incubated with a biotinylated probe against tRNA^His^_GUG_. (D) Quantitative analysis of relative levels of Q-modified tRNA^His^_GUG_. The signal intensities of the Q-tRNA band for each sample was divided by the signal intensities for their total tRNA content (Q+G) and then normalized to the wild type (T) samples. The data represent mean ± SD for three independent experiments (Unpaired Student’s t test, p = 0.0003)

These data indicate that queuine is incorporated into *E. histolytica* tRNAs, suggesting the presence of an active TGT in the parasite. The results also raise concerns about the limited amount of queuine available for *E. histolytica* in the complete TYI-S-33 medium.

Recently, it has been found that a modification of the wobble position of tRNA with a GUN anticodon by 7-deaza-guanosine derivative queuosine (Q34) stimulates *S. pombe* Dnmt2/Pmt1-dependent C_38_ methylation (m^5^C_38_) in the tRNA^Asp^_GUC_ anticodon loop [17]. The presence of an active Dnmt2 in *E. histolytica* [25–27] incited us to check if queuine also enhances the level of m^5^C_38_ in the parasite’s tRNA^Asp^_GUC_ anticodon loop (Fig 1B). We used PCR-based bisulfite sequencing to detect cytosine methylation in the tRNA by identifying the location of the methylated cytosine by sequencing. Amplicons were generated from trophozoites that were grown with and without queuine and sequences of several independent PCR products were sequenced and analyzed. We observed that the percentage of C_38_ tRNA^Asp^_GUC_ methylation in trophozoites that were grown with queuine (52.5 ± 3.5%) was significantly higher than the percentage of C_38_ tRNA^Asp^_GUC_ methylation in control trophozoites (20 ± 1.4 %). Moreover, we were unable to detect methylation in C_32_, C_33_, C_37_, C_48_, and C_56_, whereas 100% methylation was observed in C_49_. These results which are in accordance with our previously published results [25] indicate that the bisulfite reaction was successful. These results also indicate that the incorporation of queuine into amoebic tRNAs leads to hypermethylation of tRNA^Asp^_GUC_.

### Characterization of EhTGT

Given the queuine-induced increase in Q in tRNA, we next set out to characterize the *E. histolytica* TGT (EhTGT) and its role in Q incorporation and cell phenotype. We used a homology-based approach with the human QTRT1 gene to identify the subunits of *E. histolytica* TGT. The eukaryotic TGT enzymes, such as human TGT, consist of a queuine tRNA-ribosyltransferase 1 (QTRT1, eubacterial TGT homolog) and a queuine tRNA-ribosyltransferase domain-containing 1 (QTRTD1). To get more information about EhTGT, we performed a protein sequence alignment of *Homo sapiens* QTRT1 subunit (hQTRT1), EhQTRT1, and EhQTRTD1 (Fig S2A). Based on the annotation of the *E. histolytica* genome, a homolog of hQTRT1 and hQTRTD1 exists in *E. histolytica*, namely EhQTRT1 (XP_656142.1) and EhQTRTD1 (XP_652881.1). The EhQTRT1gene shares a high degree of homology with the human QTRT1 (52% sequence identity), whereas the EhQTRTD1 shares a homology at the C-terminal domain (28.3% sequence identity). Moreover, these two proteins also share a high degree of homology with the *Zymomonas mobilis* TGT (41.6% and 35.6%, respectively). The *Z. mobilis* TGT enzyme is a homodimeric zinc-binding protein, whereas the eukaryotic TGT is a heterodimeric zinc-binding protein. We identified four essential residues that are important for Zn^2+^ binding shown highlighted purple in (Cys 305, Cys307, Cys310, and His342; *E. histolytica* numbering) to be conserved in both EhQTRT1 and EhQTRTD1 (Fig S2B).

To get information about the EhTGT structure, we built an *in silico* model of EhTGT subunits based on the *Thermotoga maritima* QTRT1 structure and murine QTRT2 structures. The prediction also suggested that EhQTRT1 and EhQTRTD1 are homodimers (Fig S3 A&B), which interact together to form a heterodimer like the eukaryotic TGT enzyme (Fig 3A)[28]. To get more insights about the EhTGT enzyme, a polyhistidine-tagged EhQTRT1 and untagged EhQTRTD1 was cloned and coexpressed in *E. coli*. During protein expression, the addition of 100μM ZnS0_4_ and a low-temperature induction (19 °C) was found to be essential for obtaining an active complex. The EhQTRTD1 was copurified with the polyhistidine-tagged EhQTRT1 by Ni^2+^ affinity chromatography. The enzyme was further purified by size-exclusion chromatography (Fig 3B) and the fractions were analyzed by SDS-PAGE (Fig 3C). The molecular weight of both EhQTRT1 and EhQTRTD1 on SDS-PAGE revealed their molecular mass to be around 44.2kDa and 44.1kDa respectively, which matches their predicted molecular mass [29]. To confirm the identity of both bands, mass spectrometry was performed on peptides obtained from tryptic digests of the bands excised from the SDS gel. After mapping these peptide fragments against *E. coli* and *E. histolytica* databases, our samples were confirmed to be EhQTRT1 and EhQTRD1 (Fig S3C).

**Figure 3:**
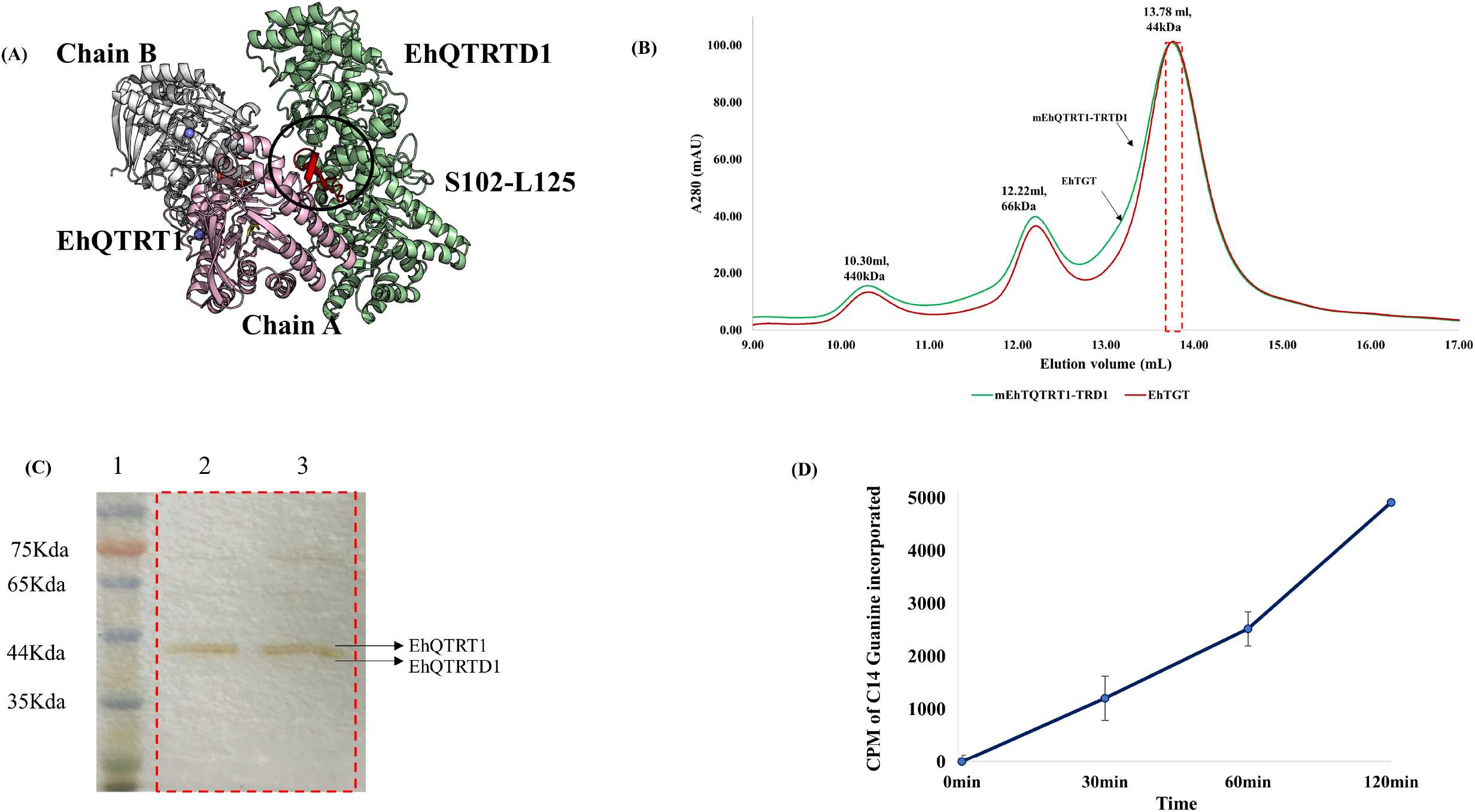
In silico modeling of EhTGT and expression in *E. coli*. (A) *In silico* model analysis showing the interaction of EhQTRT1 and EhQTRTD1 and expression of EhTGT in *E.coli*. The EhQTRT1-EhQRTD1 complex was successfully modeled. EhQTRT1 and EhQTRTD1 were co-expressed in *E.coli* using a pETDuet-1 vector. (B) A size-exclusion chromatogram representing purified EhTGT (red line) and mEhQTRT1-QTRTD1 (green line). The elution volumes of each protein have been mentioned on the top of each peak, along with their predicted molecular weight after elution from the column. (C) SDS-PAGE showing the formation of a heterodimer in EhTGT (lane 2) and mEhQTRT1-QTRTD1 (lane3). The elution volumes (shown in red box) correspond to peak obtained after size-exclusion chromatography for each protein. (D) CPM of 8-[^14^C]-guanine incorporated over time in EhtRNA^Asp^_GUC_ over time in the presence of EhTGT (blue line). The graphs represent mean ± SD of CPM counts for three independent experiments performed in triplicates.

To verify that the EhTGT is active, the activity of the complex was tested by determining its ability to displace guanine in the tRNA^Asp^_GUC_ anticodon with 8-[^14^C]-guanine. We observed that EhTGT is an active enzyme and that EhQTRT1-EhQTRTD1 enzyme complex efficiently incorporates 8-[^14^C]-guanine with a specific activity of 2.28 pmol/min/mg of protein (Table 1) (Fig 3D). However, no activity was detected when tRNA^Val^_CAC_ was used as a substrate (Table 1).

**Table 1:** Validation of the *in silico* model of EhTGT. Our results indicate that EhTGT is an active enzyme when it forms a functional complex and when tRNA^Asp^_GUC_ is used as substrate. A punctual mutation (D267 to A267) in the predicted active site of EhQTRT1 makes the enzyme (mEhQTRT1-TRD1) inactive without affecting the formation of the EhQTRT1-EhQTRTD1 complex. No EhTGT activity was detected when tRNA^Val^_CAC_ was used as substrate. The specific activity of EhTGT incubated with tRNA^Asp^_GUC_ was significantly different from mEhQTRT1-EhQTRTD1 activity and from EhTGT activity when the enzyme is incubated with tRNA^Val^_CAC_ according to an unpaired student’s t test (p≤0.05). The data are expressed as mean ± standard deviation and are representative of three independent experiments performed in duplicates.

| Enzyme | Substrate | Specific activity of the enzyme<br>(pmol/min/mg of protein) |
| --- | --- | --- |
| EhTGT | tRNA<sup>Asp</sup><sub>GUC</sub> | 2.28±0.01 |
| mEhQTRT1-TRD1 | tRNA<sup>Asp</sup><sub>GUC</sub> | 0.01±0.02 |
| EhTGT | tRNA<sup>Val</sup><sub>CAC</sub> | 0 |

The *in silico* model also predicted that the stretch of S102-L125 in EhQTRT1 is essential for the binding of EhQTRTD1, and that the D267 residue in EhQTRT1 is a substrate-binding residue (Fig 3A). A punctual mutation (D267 to A267) in the predicted active site of EhQTRT1 (indicated in a black circle in Fig 3A) made the enzyme inactive without affecting the formation of the EhQTRT1-EhQTRTD1 complex (Table 1, Fig 3C). These results confirm the prediction of the *in silico* model regarding the location of the active site (Fig 3A).

To better understand the role of EhTGT in *E. histolytica*, we silenced the expression of EhQTRT1 by using an epigenetic method based on antisense small RNAs [30]. In this method, a gene-coding region to which large numbers of antisense small RNAs map is used as a ‘trigger’ to silence the gene fused to it. The silencing is maintained after that the trigger plasmid has been cured of the silenced strain. Here, we used the siEhQTRT1 plasmid to silence the expression of EhQTRT1. Following transfection of trophozoites with the siEhQTRT1 plasmid and the selection of a population resistant to G418 (6 μg/ml), the siEhQTRT1 plasmid was cured by removal of G418 from the culture for a month. Silencing of EhQTRT1 in the cured EhQTRT1 silenced trophozoites was confirmed by immunoblotting using an antibody against EhTGT (Fig 4A) and by northern blot by using a radiolabeled probe against EhQTRT1 (Fig 4C). Using densitometry analysis, we observed that the EhTGT complex in siEhQTRT1 trophozoites was inhibited by 80% at the protein level (Fig 4B) and by 90% at the mRNA level (Fig 4D). The cellular localization of EhTGT in control trophozoites and siEhQTRT1 trophozoites has been performed by confocal immunofluorescence microscopy. In control trophozoites, EhTGT has a cytoplasmic localization which is barely detectable in siEhQTRT1 trophozoites (Fig 5A & B).

**Figure 4:**
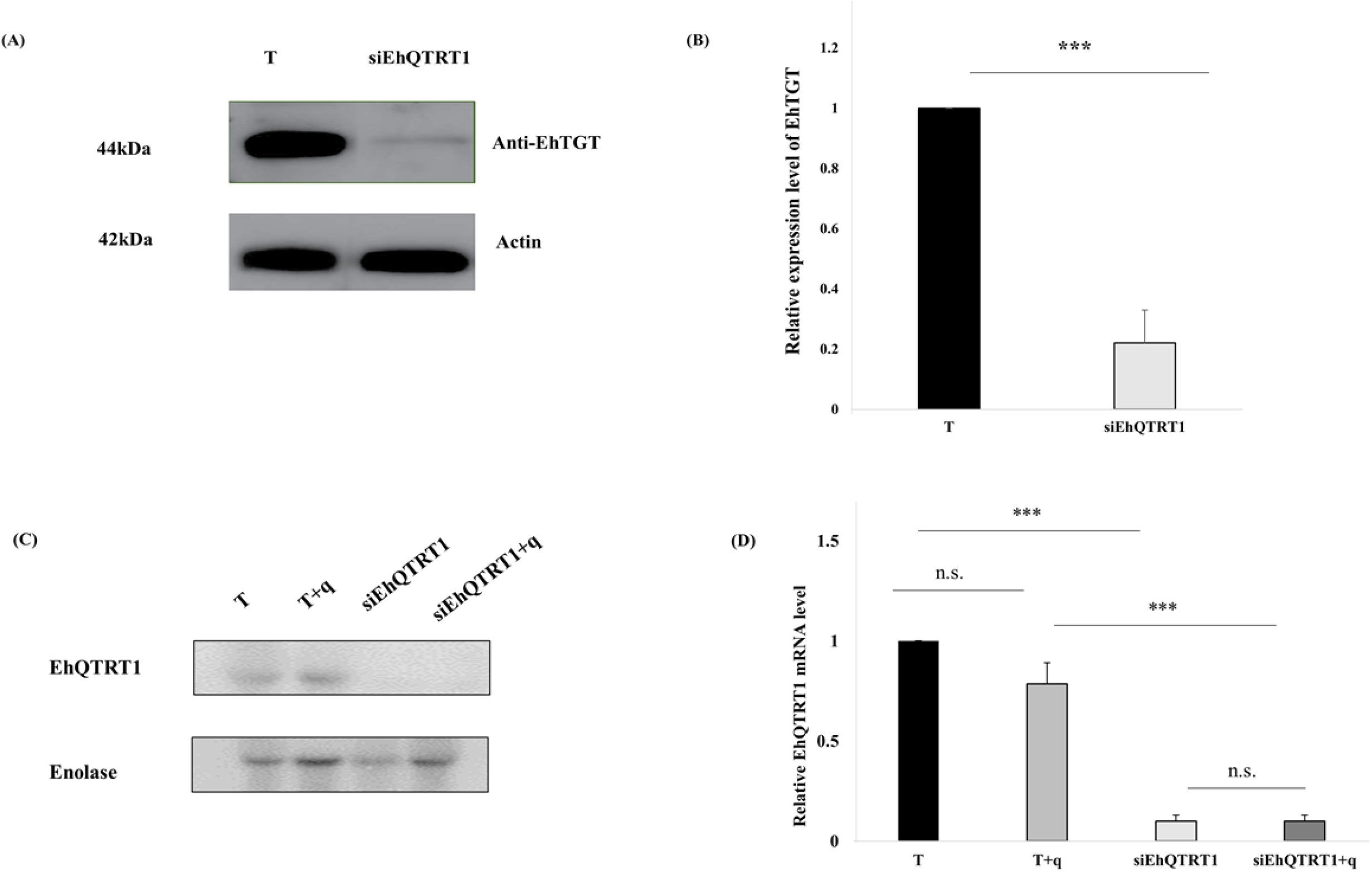
Silencing of EhTGT. (A): Western blotting was performed on total protein extracts that were prepared from wild type *E. histolytica* trophozoites and EhQTRT1 silenced trophozoites. The proteins were separated on 10% SDS-PAGE gels and analyzed by western blotting using a homemade EhTGT antibody (1:1000) or actin antibody (1:1000). (B): Densitometry analysis showing the percentage of inhibition of the EhTGT complex in EhQTRT1 silenced trophozoites (normalized to wild type). The data represent mean ± SD for three independent experiments (Unpaired Student’s t test, p = 0.0003). (C) Northern blot analysis of EhQTRT1. Enolase has been used as housekeeping gene because its expression does not change in response to queuine (this work). (D) Quantitative analysis of relative density of mRNA. The signal intensities of the samples were divided by enolase mRNA and were then normalized to the wild type mRNA. Data are represented as mean ± standard deviation about the mean for two independent experiments

**Figure 5:**
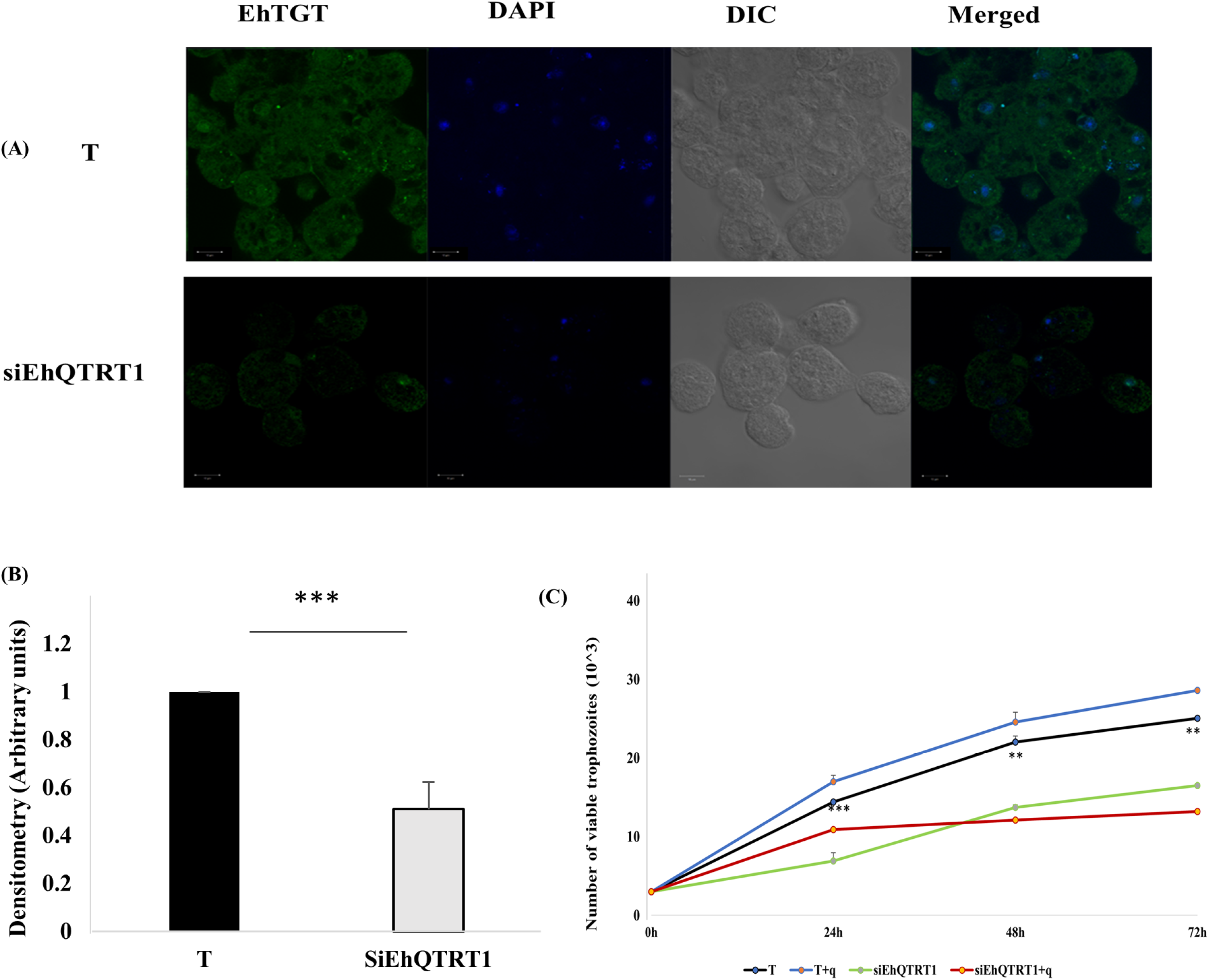
Characterization of the EhTGT silenced trophozoites. (A) EhTGT was detected in control trophozoites and siEhQTRT1 trophozoites that were grown with and without queuine using a homemade polyclonal mice EhTGT antibody (1:250) and then incubated with a goat anti mouse secondary antibody Alexa fluor (Jackson Immuno Research) at a concentration of 1:250. Nucleus are stained with 4’,6-diamidino-2-phenylingole (DAPI) (Sigma-Aldrich) at concentration of 1:1000. The samples were examined under a confocal immunofluorescence microscopy (Zeiss-LSM700 Meta laser scanning system confocal imaging system) with a 63X oil immersion objective. Images are scaled to 10μm. (B). Densitometry analysis showing the percentage of the EhTGT complex in 15 trophozoites in each condition (Normalized to control trophozoites). Data are expressed as mean ± SD representative of three independent experiments (Unpaired Student’s t test, p ≤ 0.0001). (C) Effect of silencing EhQTRT1 on the growth of *E. histolytica*. Control trophozoites (T) and Trophozoites silenced for EhQTRT1 (siEhQTRT1) (30 X10^3^ /ml of complete TYI media) were grown for 24h, 42, and 72 hours with queuine (+q) or without queuine. The number of viable trophozoites at each time point was determined by using the vital stain eosin. siEhQTRT1 trophozoites have their growth impaired compared to that of control trophozoites. Queuine has no significant effect on the growth of T or SiEhQTRT1 trophozoites. The data are representative of three independent experiments performed in duplicates. Data are expressed as mean ± SD. (Unpaired Student’s t test, T vs siEhQTRT1 24h (p= 0.0006), 48h (p= 0.0082), 72h (p= 0.005).

We have previously demonstrated that EhTGT can incorporate queuine into *E. histolytica* tRNAs (Fig 1A). Consequently, we assumed that siEhQTRT1 trophozoites will have a reduced ability to incorporate queuine in their tRNAs. To test this assumption, we have determined the global level of Q in siEhQTRT1 trophozoites tRNAs (Fig 1A) and in Q-tRNA^His^_GUG_ and tRNA^Asp^_GUC_ (Fig 2A and B). We observed a very strong reduction of the level of Q-tRNAs, Q-tRNA^His^_GUG_ and tRNA^Asp^_GUC_ in siEhQTRT1 trophozoites that were grown with queuine when compared to control trophozoites that were grown with queuine. The amount of m^5^C_38_ methylation in tRNA^Asp^_GUC_ in siEhQTRT1 trophozoites that were grown with queuine was also significantly reduced (23.5±2.1 %) compared to the level in control trophozoites that were grown with queuine (52.5±3.5%) (Fig 1B).

We also observed that siEhQTRT1 trophozoites have their growth impaired by 50% when compared to the growth rate of the control trophozoites (Fig 5C). Queuine has no significant effect on the growth of control or siEhQTRT1 trophozoites (Fig 5C).

### Queuine induces the resistance of *E. histolytica* to OS

Having established that queuine supplementation leads to increased Q levels in tRNA, we next define the effect of queuine on the parasite phenotype. To understand if queuine regulates resistance to OS in the parasite, we grew trophozoites with and without queuine and exposed them to different concentrations of H_2_0_2_ (0-5mM). We observed that the LD_50_ of H_2_0_2_ of trophozoites that were grown with queuine (3.3±0.05 mM) is significantly higher than in the absence of queuine (2.4± 0.3 mM) (Fig 6). These results indicate that queuine protects the parasite against OS. Next, we asked whether the beneficial effect that queuine has on the resistance of *E. histolytica* to OS (Fig 6) depends on EhTGT. For this purpose, we determined the resistance of siEhQTRT1 trophozoites that were grown with queuine and then exposed to OS (Fig 6). We observed that queuine does not protect siEhQTRT1 trophozoites against OS.

**Figure 6:**
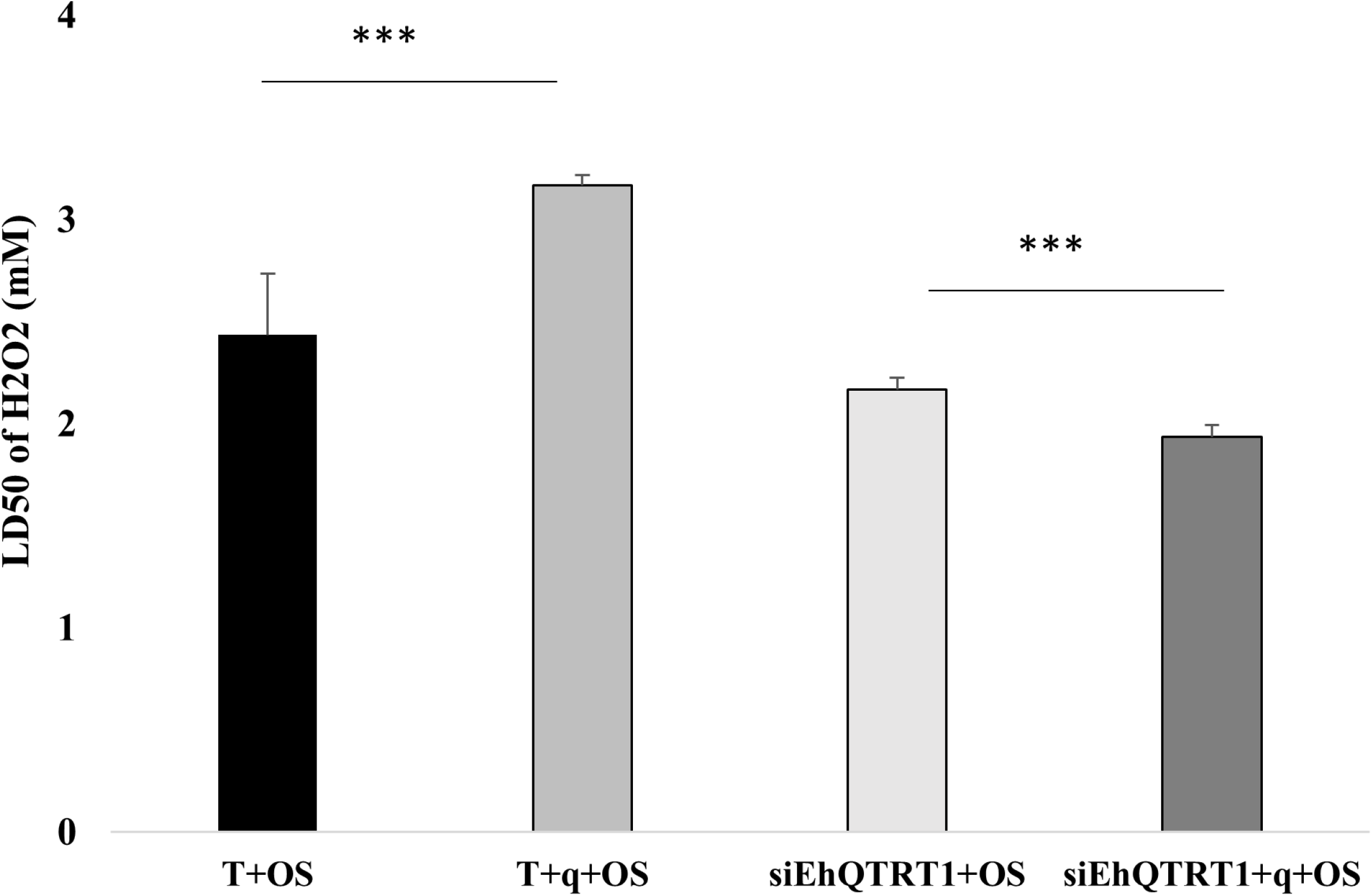
Queuine triggers resistance to OS and this effect is lost in siEhQTRT1 trophozoites. Determination of the amount of H_2_O_2_ required to kill 50% of the population (LD50) of control trophozoites, trophozoites that were grown with queuine (0.1μM for 3 days), siEhQTRT1 trophozoites and siEhQTRT1 trophozoites cultivated with queuine. The data represent mean ± SD of three independent experiments performed in duplicates. The LD_50_ of trophozoites that were grown with queuine and exposed to OS (T+q+OS) was significantly higher than the LD_50_ of untreated trophozoites exposed to OS (T) (Unpaired Student’s t test, p≤0.005). The LD_50_ of siEhQTRT1 trophozoites that were grown with queuine and exposed to OS (siEhQTRT1+q+OS) was significantly lower than the LD_50_ of siEhQTRT1 trophozoites exposed to OS (siEhQTRT1+OS) (Unpaired Student’s t test, p≤0.005)

### Queuine regulates protein synthesis during OS

The protective effect that queuine has on the parasite exposed to OS suggests that protein synthesis is preserved in the parasite exposed to queuine and OS. To test this hypothesis, we used the surface sensing of translation assay (SUnSET) [31] to determine the amount of puromycin that was incorporated into nascent peptide chains (Fig 7A&B&E). As previously described [32], we found that protein synthesis is strongly inhibited in trophozoites exposed to cycloheximide, a protein synthesis inhibitor and in trophozoites exposed to OS (Fig 7B&E). The inhibitory effect of OS on protein synthesis did not occur in trophozoites that were grown with queuine before being exposed to OS (Fig 7B&E). Interestingly, queuine seems to improve the level of protein synthesis in control trophozoites (Fig 7B&E). Next, we asked whether the regulatory effect that queuine has on protein synthesis depends on EhTGT. For this purpose, SunSET assay was performed on siEhQTRT1 trophozoites that were grown with and without queuine and then exposed or not to OS (Fig 7C&D&E). We observed that none of the effect that queuine has on protein synthesis is observed in siEhQTRT1 trophozoites.

**Figure 7:**
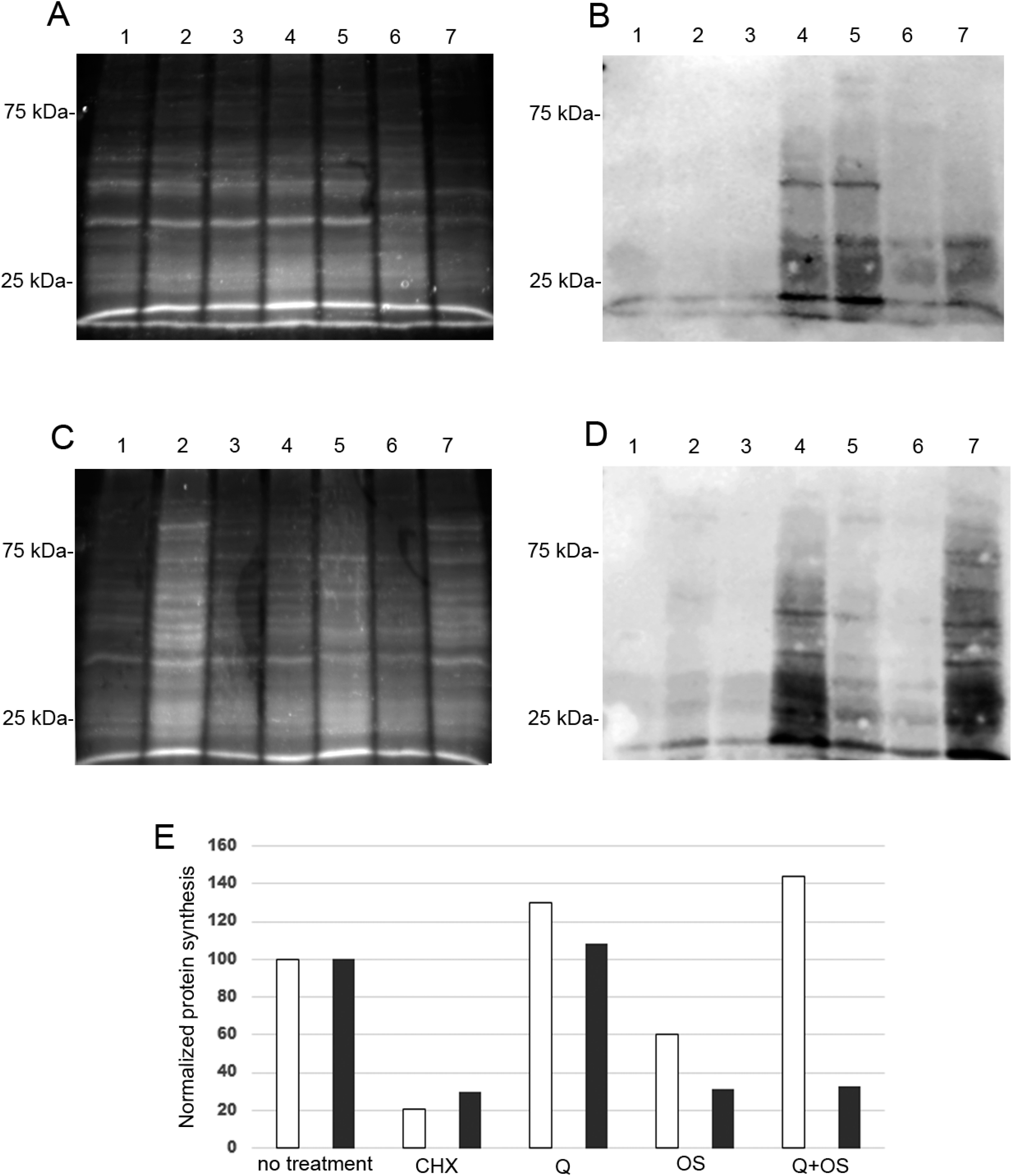
Measurement of protein synthesis in control trophozoites and in siEhQTRT1 trophozoites using puromycin-labeled proteins. (A and C) stain-free total protein labeling. (B and D) Western blot with puromycin antibody. (A and B): Lane 1-control trophozoites. Lane 2-control trophozoites that were grown with queuine (0.1μM for three days). Lane 3-control trophozoites treated with cyclohexamide (100ug/ml) before being labeled with puromycin (10μg/ml). Lane 4-control trophozoites labeled with puromycin. Lane 5-control trophozoites that were grown with queuine before being labeled with puromycin. Lane 6-control trophozoites exposed to 2.5mM H_2_0_2_ for 20 minutes before being labeled with puromycin. Lane 7-trophozoites that were grown with queuine and then exposed to 2.5mM H_2_0_2_ before being labeled with puromycin. (C and D) Lane 1-siEhQTRT1 trophozoites. Lane 2-siEhQTRT1 trophozoites that were grown with queuine (0.1μM for three days). Lane 3-siEhQTRT trophozoites treated with cyclohexamide (100ug/ml) before being labeled with puromycin (10μg/ml). Lane 4-siEhQTRT1 trophozoites labeled with puromycin. Lane 5-siEhQTRT1 trophozoites that were grown with queuine prior to their exposure to 2.5mM H202 before being labeled with puromycin. Lane 6-siEhQTRT1 trophozoites exposed to 2.5mM H202 before being labeled with puromycin. Lane 7-siEhQTRT1 trophozoites that were grown with queuine before being labeled with puromycin. E. Normalization with ImageJ of the puromycin signal (B) against the total protein signal (A) for control trophozoites (white bars) and of the puromycin signal (D) against the total protein signal (C) for siEhQTRT1 trophozoites (black bars). Normalized values for control trophozoites labeled with puromycin and for siEhQTRT1 trophozoites labeled with puromycin were taken as 100%. Total protein extracts were separated on a SDS-PAGE gel and they were analyzed by Western blotting with a mouse-monoclonal puromycin antibody (1:5000). These results are representative of two independent experiments. Total protein was determined using stain-free detection.

### Queuine impairs cytopathic activity and the survival of the parasite within the mouse caecum

It has been previously shown that TGT and presumably queuine regulates the virulence of *Shigella* by controlling the translation of VirF, a central transcriptional regulator of virulence factors involved in cellular invasion and spreading of this bacteria [33]. To test whether queuine affects the virulence of the parasite, we determined the ability of *E. histolytica* trophozoites that were grown with and without queuine to destroy a monolayer of mammalian cells (cytopathic activity) (Fig 8A). We found that the cytopathic activity of trophozoites that were grown in the presence of queuine was impaired compared to that of trophozoites that were grown in absence of queuine.

**Figure 8:**
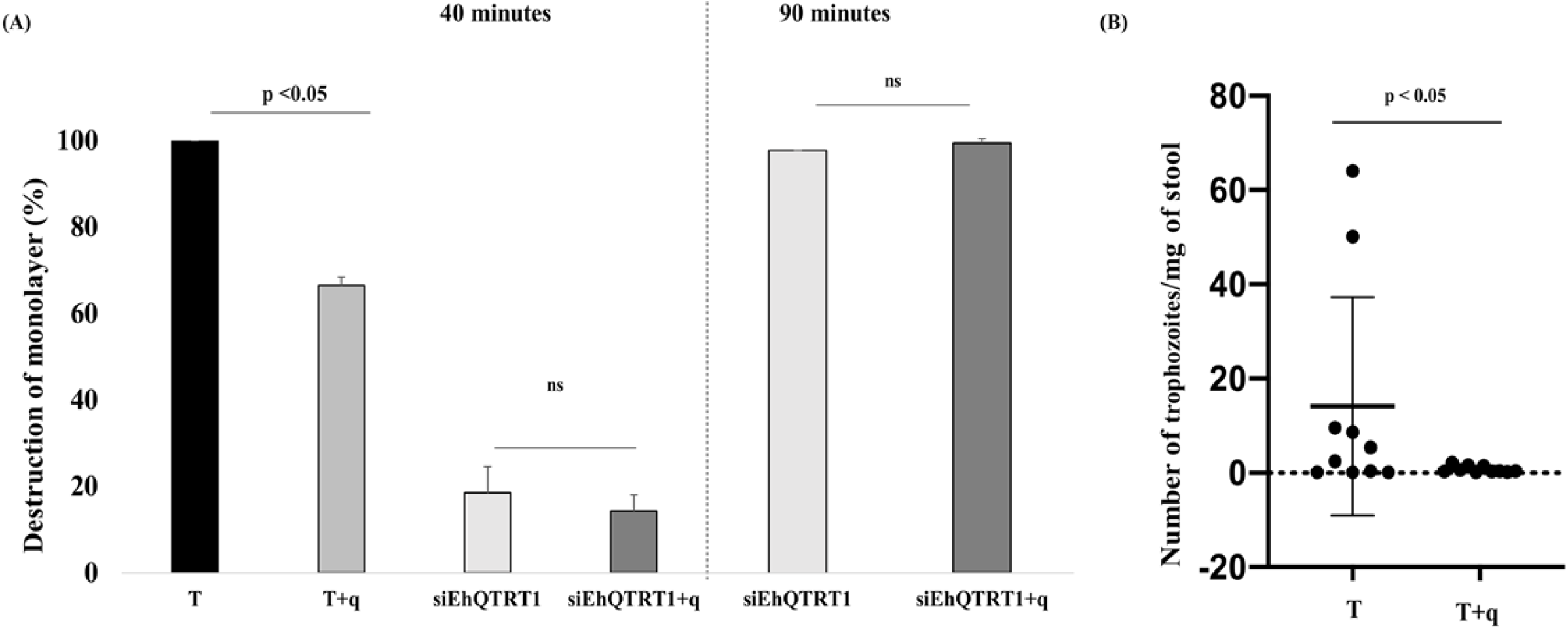
Effect of queuine on the virulence of *E. histolytica*. (A). Cytopathic activity of *E. histolytica* trophozoites and siEhQTRT1 trophozoites that were grown with and without queuine was measured by their ability to destroy a monolayer of HeLa cells. *E. histolytica* trophozoites were grown with and without queuine (0.1μM for 3 days). Data are displayed as the mean ± standard deviation of three independent experiments that were repeated twice. (Unpaired Student’s t test, T vs T+q, p= 0.0026). siEhQTRT1 trophozoites that were grown with and without queuine were incubated with HeLa cell monolayer for 40 or 90 minutes. The cytopathic activity of siEhQTRT1 trophozoites that were grown with queuine is not different from the cytopathic activity of siEhQTRT1 trophozoites that were grown without queuine at both 40 and 90 minutes of incubation between the trophozoites and the HeLa cell monolayer. The data are representative of three independent experiments (normalized to wild type) (Unpaired Student’s t test, p= 0.01 (T vs T+q)). (B) Effects of queuine treatment in intestinal amoebiasis. Number of trophozoites determined in CBA/J mice stool infected with trophozoites that were grown with queuine (q) or non-treated trophozoites (T). Trophozoites number was quantified by the quantitative real-time PCR method using *E. hisolytica* 18S rRNA specific primers. Data represent the mean ± SD for 10 mice. p < 0.05 by an Unpaired Student’s t-test.

Next, we asked whether the detrimental effect that queuine has on the cytopathic activity of *E. histolytica* (Fig 8A) depends on EhTGT. We found that the cytopathic activity of siEhQTRT1 trophozoites is significantly impaired (Fig 8A) compared to that of control trophozoites (Fig 8A). Moreover, the growth of siEhQTRT1 trophozoites in the presence of queuine did not have any significant effect on their cytopathic activity (Fig 8A). To determine if the siEhQTRT1 trophozoites require more time to destroy the HeLa monolayer than control trophozoites, we incubated siEhQTRT1 trophozoites that were grown with and without queuine on the HeLa cell monolayer for 90 minutes instead of 40 minutes. We found that more than 90 % of the HeLa monolayer is destroyed by siEhQTRT1 trophozoites after 90 minutes of incubation (Fig 8A). The same level of monolayer destruction was observed for siEhQTRT1 trophozoites that were grown with queuine (Fig 8A). These results indicate that queuine impairs the strong cytopathic activity of control trophozoites (Fig 8A) but does not impair the weak cytopathic activity of siEhQTRT1 trophozoites.

To validate the alteration of amoebic pathogenicity *in vivo*, trophozoites that were grown in the presence of queuine were subjected to the mouse intestinal amoebiasis (Fig 8B). In mouse intestinal amebiasis model, control trophozoites or trophozoites that were grown in the presence of queuine were inoculated to susceptible CBA/J mice cecum. After 7 days of post-inoculation, the number of trophozoites present in the stool was determined. The average number of detectable trophozoites from mice inoculated with control trophozoites (14 ± 23 trophozoites/mg stool) was higher than that of trophozoites that were grown in the presence of queuine (0.74 ± 0.71 trophozoites/mg stool) (Fig 8B). These results indicate that queuine affects the survival of the parasite in the mouse caecum.

### Queuine has a profound impact on the *E. histolytica* transcriptome

We used RNA sequencing (RNA-seq) to investigate the molecular basis of the effect of queuine on the resistance of *E. histolytica* to OS (Fig 9A and B). RNA-Seq experiments were performed as described previously [10], and the data (available at http://www.ncbi.nlm.nih.gov/geo under the accession number GSE142211) were analyzed using the generalized linear model implemented in the DESeq2 package in R. This allowed us to perform pairwise comparisons of gene expression by the *E. histolytica* HM1:IMSS strain under the various tested conditions and to probe the interaction between queuine and OS (Fig. 9B).

**Figure 9:**
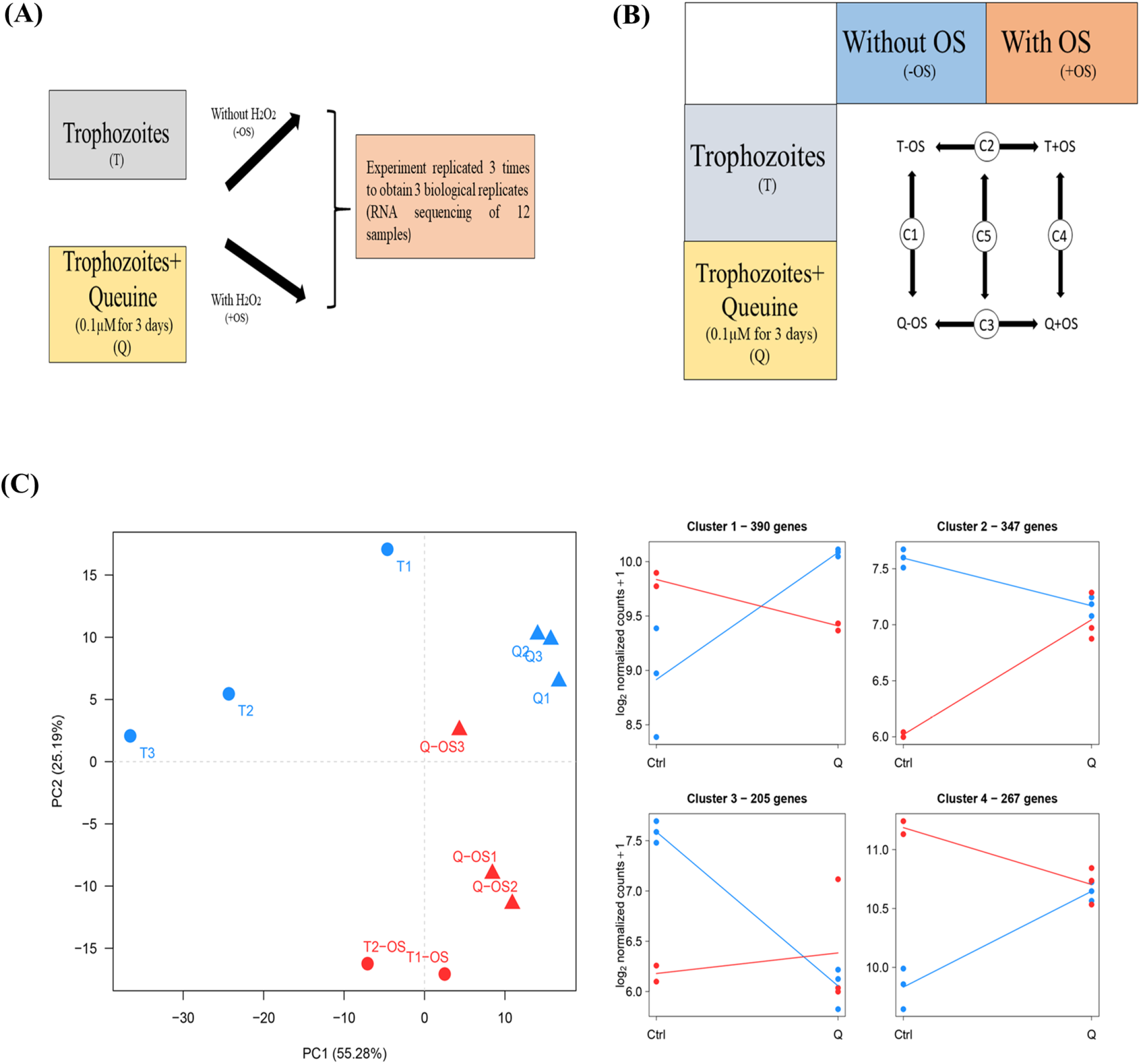
Experimental design of the RNA-seq. (A) Trophozoites have been grown with and without 0.1μM queuine for 3 days (T and q), and then were exposed (or not) to OS (+OS and −OS). The experiments were repeated three times to obtain 12 samples that were purified and sequenced further. (B). Biological conditions compared during differential RNA-seq analysis. 5 pair-wise comparisons have been performed. (C) Principal Component Analysis of the RNA-Seq data performed on the variance-stabilizing transformed count matrix. Average profiles for the four gene clusters defined by the hierarchical clustering. The average transcriptome profile was established by computing the mean log_2_-normalized count for the genes in each of the four identified clusters.

We first used a principal component analysis to explore the data’s structure (Fig. 9C); this showed that the untreated trophozoites group (T) and the trophozoites that were grown with queuine (Q) form two distinct clusters that can be separated on the first principal component. Stress effect is mainly supported by the second component and OS-treated and -untreated samples are also forming clusters both in T and in Q conditions. Our comparisons revealed that queuine has a profound impact on the *E. histolytica* transcriptome (C1) with 664 downregulated and 548 upregulated genes (Table 2). Details regarding these genes can be found in Table S1.

**Table 2:** Number of genes modulated in *E. histolytica* trophozoites in control trophozoites (T) and in trophozoites that were grown with queuine (q) in response to OS.

|  | Comparison | Modulated Genes |  |  |
| --- | --- | --- | --- | --- |
|  |  | # Downregulated | # Upregulated | Total |
| C1 | Q-OS vs T-OS | 664 | 548 | 1212 |
| C2 | T+OS vs T-OS | 985 | 661 | 1646 |
| C3 | Q+OS vs Q-OS | 248 | 92 | 340 |
| C4 | Q+OS vs T+OS | 176 | 265 | 441 |
| C5 | C3 vs C2 | 485 | 447 | 932 |

### Gene categories affected by queuine supplementation

The differentially regulated genes in Q vs T (comparison C1) were classified according to the protein class which they encode (Fig 10A&B) using the Protein ANalysis THrough Evolutionary Relationship (PANTHER) sequence classification tool (version 14.1)[34]. Of the upregulated genes in the presence of queuine (Fig 10A), genes that are associated with nucleic acid binding (GO:0003676) and nucleosome binding (GO:0031491) such as chromatin-specific transcription elongation factor (EHI_109860) and protein HIRA (EHI_131950), heterocyclic compound binding (GO:1901363) such as 60S ribosomal protein L4 (EHI_064710) and ATP-dependent DNA helicase (EHI_119290), ATPase activity (GO:0016887) such as heat shock protein 70 family (EHI_001950) and RecQ family helicase (EHI_028890), oxidoreductase activity (GO:0016491) such as peroxiredoxin (EHI_145840) and aldehyde-alcohol dehydrogenase 2 (EHI_024240), catalytic activity, acting on RNA (GO:0140098) such as regulator of nonsense transcripts (EHI_035550) and RNA lariat debranching enzyme (EHI_062730), actin binding (GO:0003779) such as coronin (EHI_122800) and villidin (EHI_150430), cytoskeletal protein binding (GO:0008092) and tubulin binding (GO:0015631) such as dynamin-1-like protein (EHI_013180) and kinesin-like protein (EHI_140230), protein-containing complex binding (GO:0044877) such as elongation factor 2 (EHI_189490) and chromatin-specific transcription elongation factor (EHI_109860), catalytic activity, acting on DNA (GO:0140097) such as DNA replication licensing factor (EHI_076880) and RecQ family helicase EHI_028890 are significantly enriched according to the PANTHER statistical overrepresentation test (Fig 10C).

**Figure 10:**
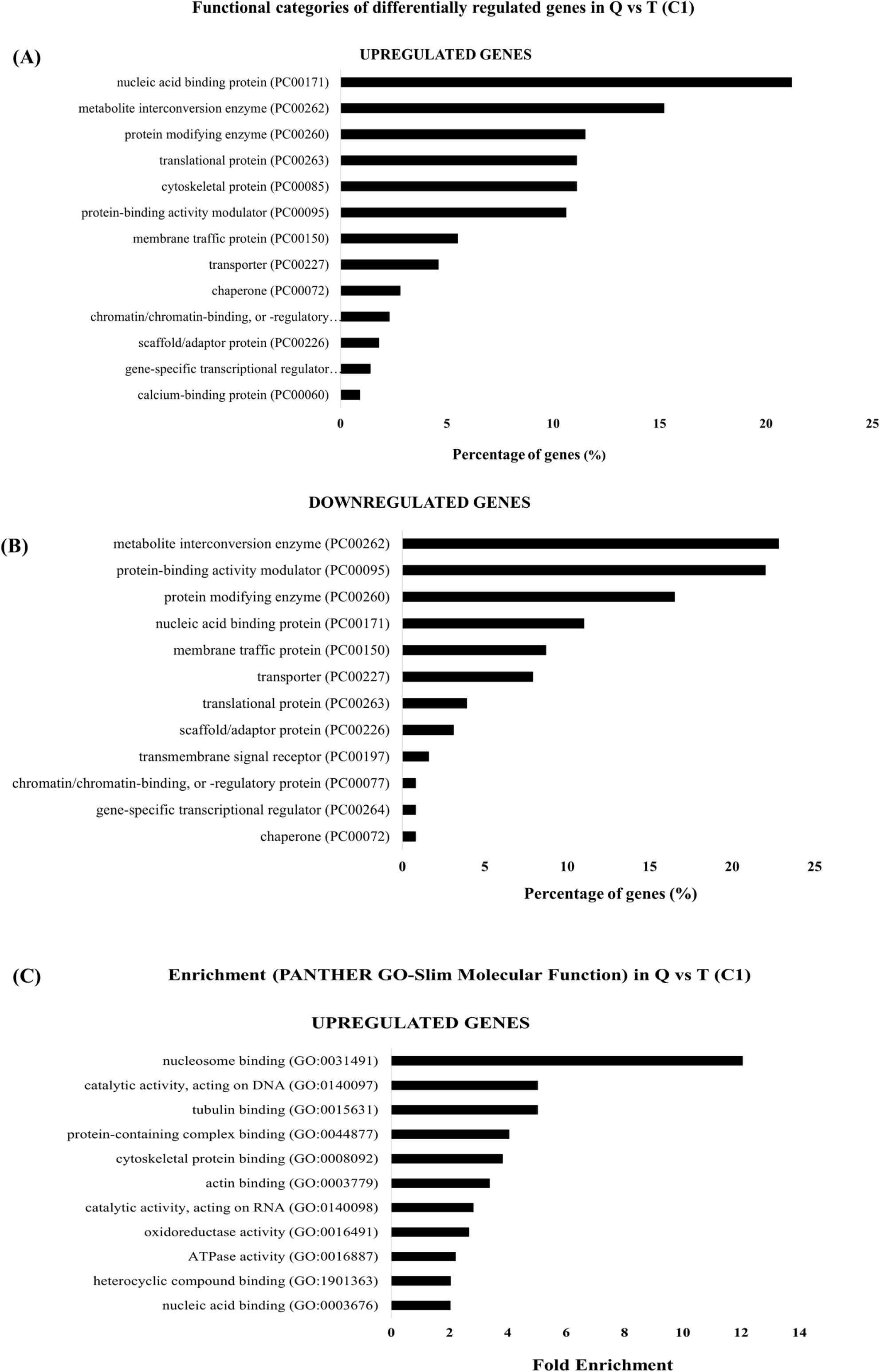
Functional categories and classification of genes enriched in trophozoites that were grown with queuine vs control trophozoites. Functional categories: **(A)** up regulated genes in trophozoites that were grown with queuine. **(B)** downregulated genes in trophozoites that were grown with queuine. The number of genes related to each category was derived by processing the data (Table S1) with PANTHER (http://pantherdb.org). **(C)** Enrichment test in queuine treated trophozoites. The details of genes present in each cluster are given in Table S3.

The categories for functional classification of genes downregulated in trophozoites that were grown with queuine (using the subset of protein class) is shown in (Fig 10B). The most abundant class of proteins are metabolite interconversion enzyme (PC00262) (22.8%), protein-binding activity modulators (PC00095) (22%), Protein modifying enzymes (PC00260) (16.5%), nucleic acid binding protein (PC00171) (PC00262) (11%), membrane traffic protein (PC00150) (8.7%), transporter (PC00227) (7.9%), translational protein (PC00263) (3.9%), scaffold/adaptor protein (PC00226) (3.1%).

Of the downregulated genes in Q, no enrichment of a specific biological process was detected according to the PANTHER statistical overrepresentation test (Fig 10B).

### Gene categories affected by queuine supplementation and OS

As previously described [10], OS strongly modulates the transcriptome of the parasite with 1646 genes that are differentially expressed compared to the T condition (comparison C2). Interestingly, in the presence of queuine, OS affects the expression of only 340 genes, suggesting that queuine influences the response of the parasite to OS. Next, we addressed the influence of queuine on *E. histolytica*’s response to OS (comparison C5). We identified 932 genes (Table 2), and then performed a hierarchical clustering analysis (Fig. 9C). The average profile of each of the four identified clusters was then established (Fig. 9C) (Table S3). Clusters 1&4 represent genes that are mostly upregulated in trophozoites exposed to OS but that are either mostly not differentially expressed (cluster 4) or down-regulated (cluster 1) in trophozoites that were grown with queuine and exposed to OS. Clusters 2&3 represent genes that are mostly downregulated in control trophozoites exposed to OS but that are mostly not differentially expressed in trophozoites that were grown with queuine exposed to OS. A Detailed analysis of these clusters is described below.

### Hierarchical clustering analysis

Cluster 1 comprised of 390 genes. 60.5% of these genes have their expression upregulated in control trophozoites exposed to OS vs trophozoites non-exposed to OS. In trophozoites that were grown with queuine exposed to OS vs trophozoites that were grown with queuine non-exposed to OS, 48% of the genes present in cluster 1 have their expression downregulated and 52% are not differentially expressed (Table 3).

**Table 3:** Summary of the hierarchical clustering analysis

|  | Comparison<br>T+OS vs T-OS |  |  | Comparison<br>Q+OS vs Q-OS |  |  |
| --- | --- | --- | --- | --- | --- | --- |
|  | Up | Down | Not DE | Up | Down | Not DE |
| **Cluster1 (390)** | 236<br>(60.5%) | 2<br>(0.51%) | 152<br>(38.97%) | 0<br>(0%) | 186<br>(47.69 %) | 204<br>(52.31 %) |
| **Cluster 2 (347)** | 0<br>(0%) | 329<br>(95%) | 18<br>(5.2%) | 9<br>(2.6%) | 23<br>(6.6%) | 315<br>(91%) |
| **Cluster 3 (205)** | 0<br>(0%) | 169<br>(82.4%) | 36<br>(17.5%) | 27<br>(13.1%) | 0<br>(0%) | 178<br>(87%) |
| **Cluster 4 (267)** | 265<br>(99.2%) | 0<br>(0%) | 2<br>(0.75%) | 28<br>(10.49%) | 11<br>(4.12%) | 227<br>(85%) |

Cluster 2 comprised 347 genes; 95% of these genes have their expression downregulated in control trophozoites exposed to OS vs trophozoites non-exposed to OS. In trophozoites that were grown with queuine exposed to OS vs trophozoites that were grown with queuine non-exposed to OS, 91% of the genes present in cluster 2 have their expression not differentially expressed. (Table 3)

Cluster 3 comprised 205 genes; 82.5% of these genes have their expression downregulated in control trophozoites exposed to OS vs trophozoites non-exposed to OS. In trophozoites that were grown with queuine exposed to OS vs trophozoites that were grown with queuine non-exposed to OS, 87% of the genes present in cluster 3 have their expression not differentially expressed. (Table 3)

Lastly, 99% of the 267 genes in cluster 4 were upregulated in control trophozoites exposed to OS vs trophozoites non-exposed to OS and 85% were not differentially expressed in trophozoites that were grown with queuine exposed to OS vs trophozoites that were grown with queuine non-exposed to OS. (Table 3)

### Functional categories identified in queuine and OS treatments

The analytical tools in PANTHER (http://pantherdb.org/) were used for functional classification and gene set overrepresentation tests. To identify the functional categories, we used the PANTHER tools, which includes entries from *E. histolytica* for functional classification. The categories for functional classification within each cluster (using the subset of protein class) are shown in Fig S4 and Table S4. The most abundant class of proteins in cluster 1 are nucleic acid binding (PC00171) (20.3%), enzyme modulator (PC00095) (16.4%) and hydrolase (PC00121) (16.4%) (Fig S4A).

The most abundant class of proteins in cluster 2 are Enzyme modulator (PC00095) (22%), Hydrolase (PC00121) (17%), and nucleic acid binding (PC00171) (14%) (Fig S4B).

The most abundant class of protein in cluster 3 are nucleic acid binding (PC00171) (21%), hydrolase (PC00121) (19%) and transferase (PC00220) (15%) (Fig S4C).

Finally, the most abundant class of protein in cluster 4 are nucleic acid binding (PC00171) (20.3%), hydrolase (PC00121) (19%) and enzyme modulator (PC00095) (13%) (Fig S4D).

### Enrichment Molecular functions

We restricted our analyses to the families of factors enriched more than 2.0-fold in the overrepresentation test (Fig S5) (Table S4).

### Functional analyses of genes in cluster 1

The bioinformatics analysis of genes presents in cluster 1 (Fig S5A) showed significant enrichment in 18 GO terms; It highlighted genes encoding proteins related to lipid transport (steroid-binding activity (GO:0005496) fold enrichment (FE):15.4 and lipid transporter activity (GO:0005319) FE: 5.7 including oxysterol binding protein and Phospholipid-transporting ATPase). It also highlighted genes encoding proteins related to catalytic activity on DNA and RNA (GO:140097 FE:4.6 and GO:140098 FE:3.7 including DNA ligase, ATP-dependent RNA helicase, recQ family DNA helicase and tRNA-dihydrouridine synthetase and tRNA synthetase (Arg, Leu, Ala, Lys, Thr, Ile, Glut, His). Cluster 1 was also enriched in genes encoding a cytoskeletal binding protein (GO:0008092) (FE:4.1) including Kinesin, Dynamin, Filamin and Villidin and Heat shock protein binding (GO:0031072) (FE:3.4) including Heat shock protein 70.

### Functional analyses of genes in cluster 2

The bioinformatics analysis of genes presents in cluster 2-showed significant enrichment in six GO terms all related to small GTPase that appeared in several combinations in the GO term enrichment analysis (Fig S5B). These include nucleoside triphosphate regulator activity (GO:0060589) (FE:3.3), GTPase regulator activity (GO:0030695) (FE: 3.45), GTPase regulator activity (GO:005096) (FE: 3.49), Ras GTPase binding (GO:0017016) (FE: 4.69), small GTPase biding (GO:0031267) (FE: 4.69), Rab GTPase binding (GO:0017137) (FE: 5.2).

### Functional analyses of genes in cluster 3

The bioinformatics analysis of genes presents in cluster 3 did not show significant enrichment.

### Functional analyses of genes in cluster 4

The bioinformatics analysis of genes presents in cluster 4 showed significant enrichment in 5 GO terms related to ribosomal proteins that appeared in several combinations in the GO term enrichment analysis (Fig S5C). These include ribosome binding (GO:0043022) (FE: 15.6), ribonucleoprotein complex binding (GO:0043021) (FE: 9.1), protein-containing complex binding (GO:0044877) (FE: 6.09), structural constituent of ribosome (GO:0003735) (FE: 4.0), and structural molecule activity (GO:0005198) (FE: 3.38).

### Comparative analysis of genes upregulated in virulent *E. histolytica* trophozoites vs genes downregulated in trophozoites that were grown with queuine

We compared our RNA Seq data containing genes downregulated in trophozoites that were grown with queuine with the previously published RNA Seq data of genes upregulated in a highly virulent *E. histolytica* HM1:IMSS strain [35]. Our analysis showed that out of 816 upregulated genes present in the virulent strain vs 664 genes downregulated in trophozoites that were grown with queuine, a total of 70 genes were common between both sets (Table S2) These genes include putative cysteine protease (EHI_151440), putative thioredoxin (EHI_152600), (EHI_124400), fatty acid elongase (EHI_009370), ubiquitin putative (EHI_156660), putative eukaryotic elongation factors (EHI_006170), and a majority of the genes correspond to hypothetical proteins that are shared between the two data sets.

Functional analysis of the proteins in this set showed that the most abundant class of proteins are protein modifying enzyme (PC00260) (PC00171) (30%), nucleic acid binding protein (PC00171) (20%), translational protein (PC00263) (20%), transporter (PC00227) (10%), protein-binding activity modulator (PC00095) (10%), metabolite interconversion enzyme (PC00262) (10 %) (Table S2).

Taken together, this transcriptome analysis indicates that queuine induces the upregulation of genes involved in OS stress response and the downregulation of genes previously associated with virulence.

### Codon usage frequency in genes which are upregulated or downregulated in trophozoites cultivated in presence of queuine

In order to gain insights into the role of Q-tRNA modification on translation in *E. histolytica*, we compared the Gene Specific Codon Usage (GSCU) frequency of each transcript upregulated in the parasite cultivated in the presence of queuine to the average codon usage frequency defined by the entire set of *E. histolytica* genes (Table S5A). Previously GSCU analysis has been used to analyze all genes in a genome to identify those overusing specific codons translationally enhanced by wobble U modifications [36] and here we have advanced the methodology to analyze transcriptionally regulated transcripts (Table S5B). We observed that some groups of genes have a distinct codon bias but we were not able to link these gene clusters back specifically to codons decoded by Q (GUN anticodon) (Fig S6A). Most likely the use of all 62 codons in our analysis obscures the clustering, as strong biases in many codons can be observed for a group of queuine regulated genes. We next stratified the transcripts upregulated in the parasite cultivated in presence of queuine based on whether they overuse the four U (U-GUN) or four C (C-GUN) ending codons decoded by Q (Table S5C). We observed two distinct groups that were classified according to the protein class which they encode (Fig S7). Of the genes that overuse the C (C-GUN), genes encoding translational protein (PC00263) such as Elongation factor 2 (EHI_166810; EHI_189490; EHI_164510) and numerous ribosomal proteins EHI_093580EHI_161180EHI_199990EHI_055670) are the most abundant (Fig S7A). Of the genes that overuse the U (U-GUN), genes encoding proteins associated with nucleic acid binding (PC00171) such as transcription initiation factor IIIB (EHI_158020), CCR4/NOT transcription complex subunit 3 (EHI_119550) or the zinc finger protein (EHI_055640) are the most abundant (Fig S7B). They are followed by genes encoding metabolite interconversion enzyme (PC00262) such as NAD FAD-dependent dehydrogenase (EHI_099700), peroxiredoxin (EHI_001420) and NADP-dependent alcohol dehydrogenase (EHI_023110) (Fig S7B).

## Discussion

We have recently demonstrated that *E. histolytica* uses oxaloacetate produced by *E.coli* to protect itself against OS [9]. In this study, we present another facet of this protective effect through the lens of the human microbiome. Queuine released by the gut bacteria has a protective effect on the parasite during OS. Interestingly, this protective effect against OS conferred by queuine is also found in cancer cells where the activity of antioxidant enzymes is improved by the supplementation of queuine in the culture [20], suggesting that this effect is universal.

In this work, we have shown for the first time that *E. histolytica* has an active TGT enzyme. EhTGT activity is essential for the incorporation of queuine in tRNAs and the resistance to OS. EhTGT shares structural similarity with its eukaryotic counterpart which is also a heterodimer. It has been proposed that one subunit (QTRT1) is responsible for the recognition of the anticodon loop of the tRNA and transglycosylase activity, whereas the second subunit (QTRTD1) is responsible to maintain the orientation of the tRNA [37, 38]. It has been suggested that the human QTRTD1 salvage queuine from queuosine-5’-monophosphate that is generated after tRNA turnover [39]. Since the amount of queuine which circulates in the range of 1-10 nM [23], it might be possible that in *E. histolytica*, there is a requirement for a second subunit to salvage Q from microbial tRNAs. Alternatively, the parasite may rely on a dedicated enzymatic machinery to salvage Q. DUF2419 is a protein with structural similarity with DNA glycosidases that has been involved in Q salvage in *S. pombe* [40]. *E. histolytica* expresses a gene, EHI_098190, which is strongly homolog to *S. pombe* DUF2419 (query cover 97%; E value 1E-28; percentage identity 27.1%). Work is in progress to characterize the involvement of EhDUF2419 in the salvage of Q from bacteria.

Different mechanisms can explain why queuine protects *E. histolytica* against OS. The first mechanism described in *S. pombe* and mammals involves the stimulation of Dnmt2 activity by prior queuosine incorporation at G34 of tRNA^Asp^_GUC_ [41, 42]. Dnmt2 mediated C38 methylation of tRNA^Asp^_GUC_ during OS protects it against cleavage by the RNA endonuclease angiogenin [43, 44]. In *E. histolytica*, we have also correlated the hypermethylation of tRNA^Asp^_GUC_ catalyzed by the *E. histolytica* Dnmt2 homolog Ehmeth with the resistance of the parasite to OS [18]. Then, this specific tRNA^Asp^_GUC_ is preferentially used by the cells for the translation of stress proteins [25]. In our current study, we have shown that exogenous supplementation of *E. histolytica* trophozoites with queuine leads to hypermethylation of C38 in tRNA^Asp^_GUC_ and that this hypermethylation depends on the active EhTGT enzyme. Thus, by extrapolating this information, it is possible that queuine exerts a protective effect on *E. histolytica* during OS via the hypermethylation of C38 in tRNA^Asp^_GUC_. The fact that EhTGT has a cytoplasmic localization (this work) whereas Ehmeth has a nuclear localization [45] does not rule out this possibility because tRNAs are able to shuttle between the cytoplasm and the nucleus [46].

In *S. pombe*, Q-tRNA modification enhances the translational speed of the C-ending codons for Asp (GAC) and histidine (CAC) and reduces that of U-ending codons for asparagine (AAU) and tyrosine (UAU) [47]. In human cells, Q-tRNA modification has also an effect on translational speed but contrary to *S. pombe*, it increases translational speed of the U-ending codons [48]. Although we did not determine in this study the effect of Q-tRNA modification on the translational speed of C vs. U ending codons, we were able to identify among the genes upregulated in the parasite cultivated in presence of queuine, two groups that either overuse the four U (U-GUN) or the four C (C-GUN) ending codons decoded by Q. Remarkably, the most abundant groups that overuses the U (U-GUN) ending codons includes possible transcription factors and proteins involved in OS resistance[49–51]. In contrast, the most abundant group that overuses the C (C-GUN) ending codons includes proteins involved in translation. In regards to the effect that queuine has on regulating gene expression, increasing resistance to OS and reducing translation in *E. histolytica*, it is tempting to speculate that U (U-GUN) ending codons will be associated with preferentially translated proteins whereas C (C-GUN) ending codons will be associated with proteins with reduced translation. Further translational, RNA modification and codon analytic studies will be used in the future to test these predictions.

The second mechanism of protection against OS mediated by queuine relies on its ability to prepare the parasite to resist OS. We have reported here that several genes involved in stress response are upregulated in the presence of queuine including heat shock protein 70 (Hsp 70), antioxidant enzymes, and enzymes involved in DNA repair. Hsp70 is known for its role in the refolding of denatured and misfolded proteins, and translocation of secretory proteins [52]. In *E. histolytica*, Hsp70 is produced by cells in response to OS, heat shock response, and their levels are also increased during the formation of liver abscess [53–55]. Antioxidant enzymes such as alcohol dehydrogenases (ADH2) are well known for their role in energy metabolism in *E. histolytica* during OS [56] and help the parasite to resist OS [57, 58]. Transketolase is another antioxidant enzyme that is involved in the pentose phosphate pathway. This enzyme is essential for the production of the antioxidant NADPH which ultimately protects cancer cells against OS stress [59, 60]. The upregulation of the expression of this enzyme in the parasite exposed to queuine (this work) and the accumulation of sedoheptulose-7-phosphate, the product of transketolase, in the parasite exposed to OS [61] suggest that this enzyme is also involved in the resistance of the parasite to OS. RecQ helicases are conserved across all forms of life ranging from prokaryotes to eukaryotes [62]. They have many metabolic functions in the cell but are widely popular for their role in DNA repair and genome stability [63–65]. Mutations in these enzymes lead to chromosomal instability which in turn is associated with many diseases such as cancer [66]. Human RecQ helicase is involved in the repair of oxidative DNA damage [66, 67], and it interacts with Poly[ADP-ribose]polymerase1 (PARP-1) both *in vivo* and *in vitro* aiding in the repair [68]. This interaction with PARP-1 is interesting as PARP-1 itself is known for its involvement in DNA repair and also in the regulation of transcription in the cell [69]. Based on these observations, we hypothesize that either queuine stimulates the production of DNA repairing enzymes in *E. histolytica* under normal physiological conditions or it prepares the parasite to resist DNA damage in response to stress. These results suggest that queuine has a role in DNA repair damage in the parasite, however, this assumption needs thorough investigation.

In this work, we have shown that queuine impairs the cytopathic activity of the parasite and its survival in the large intestine of mice with experimentally induced amebiasis. The transcriptome of the parasite exposed to queuine provides some insights into the mechanism that leads to this low virulence phenotype. Many protein modifying enzymes such as cysteine proteases, metalloproteases, phosphatases, and ubiquitin ligases have their expression downregulated in the presence of queuine. Protein phosphatases have an essential role in cellular signaling pathways and it may also play a role in the pathogenicity of the parasite [70]. GTPases are an essential part of cell signaling events and we found many genes belonging to putative Ras GTPases, Rho GTPase, and Rab GTPase proteins. Many of these genes have been linked to vesicular trafficking. For example, Rab GTPases may play an essential role in phagocytosis [71]. Moreover, a Rab GTPase in *E. histolytica* (EhRab7A) is involved in the transport of cysteine protease to phagosomes [72]. Further investigation regarding these enzymes will help us define their role in the virulence of the parasite. The most surprising observation in genes downregulated in response to queuine is the presence of 760 genes encoding hypothetical proteins. These proteins whose functions are unknown may probably play important roles in regulating the virulence of the parasite.

Another explanation as to how or why queuine may affect the virulence of the parasite stems from our comparative analysis of genes upregulated in the virulent strain [35] that are downregulated in the presence of queuine. For example, actin which is a major cytoskeletal protein involved in motility of the parasite and phagocytosis [73] has its expression upregulated in the virulent strain of *E. histolytica* and downregulated in the parasite exposed to queuine. The same trend is observed for cysteine proteases, which are essential virulence factors in the parasite that promote intestinal invasion by degrading the extracellular matrix [74]. Among the 70 common genes, more than half correspond to hypothetical proteins. Characterization of these genes presents an opportunity to discover their novel functions in the parasite. Moreover, new hypothetical proteins may serve as good markers and act as possible pharmacological targets against this parasite.

Our results may provide some clues to understand why most of the infections with *E. histolytica* are asymptomatic. Factors leading to such outcome are the level of virulence of the parasite, the immune status of the host and most probably the gut microbiota. Different mechanisms may influence the development of amoebiasis by the gut microbiota including competition with the parasite to invade the colonic mucosa or the boosting of a host immune response against the parasite [75, 76]. Here, we are adding a new mechanism to this list that is based on the downregulation of virulence genes expression by queuine. The fact that queuine confers resistance to OS seems counterintuitive because this resistance often correlates with a stronger virulence of the parasite [50, 77]. However, the resistance to OS conferred by queuine is mild if we compare it to the strong resistance conferred by oxaloacetate, an antioxidant metabolite scavenged by the parasite from the gut microbiota [78]. This mild resistance to OS may be enough to help the parasite survive fluctuation of the oxygen content that occurs in the large intestine due to changes in the composition of the microbiota. For example, the depletion of butyrate-producing species following an antibiotic treatment increased epithelial oxygenation and allows the expansion of aerobic luminal microbes [79]. The price paid by the parasite to acquire a mild resistance to OS is a lower virulence. The concept of fitness cost is well known and it has been described for many microorganisms [80] that have acquired drug resistance including in *E. histolytica* [81].

In this work, we have shown that queuine improves protein translation in control trophozoites and in trophozoites exposed to OS. Queuine can act at the level of translational speed, translational accuracy, protection of Q-tRNA against cleavage and amynoacylation [42]. Our attempts to separate aminoacylated from non-aminoacylated tRNA^Asp^_GUC_ or tRNA^His^_GUG_ by using acid urea polyacrylamide gel [82] were not successful. Consequently, we cannot test the effect that queuine has on tRNA aminoacylation. In *E. histolytica* and in other systems [83], stress is partially regulated by the phosphorylation of the alpha subunit of eukaryotic initiation factor 2 (eIF2) that leads to the inhibition of eIF2α activity resulting in a general decline in protein synthesis. It is possible that queuine reduces the phosphorylation level of eIF2α and consequently improve protein synthesis. Work is in progress to test this hypothesis.

Queuine regulates the transcription of more than 1200 genes. A different hypothesis can explain the effect of Q on the transcriptome of *E. histolytica:* (i) The presence of queuosine in the anticodon loop position 34 of tRNA^Asp^_GUC_ and maybe other tRNAs influences the transcriptome of *E. histolytica*.

This hypothesis is supported by a recent report showing that the deletion of a gene encoding for KEOPS (an enzyme involved in the catalysis of t6A modification in tRNA) in *S.cerevisiae* leads to the upregulation of the transcription of genes related to mitochondrial function and carbohydrate metabolism [84]. The effect of tRNA modifications on transcription can also be indirect when it impairs the translation Gcn4, a transcription factor [84]. It is possible that like for Gcn4, a yet to be discovered transcription factor whose translation depends on the presence of Q-tRNAs is involved in the regulation of gene expression in Q-trophozoites. It is also possible that Q is regulating transcription by using a mechanism that does not involve the modification of tRNA. An alternative explanation is that queuine regulates the expression of a transcription factor that controls the expression of these genes. The best-studied example is in the enteropathogenic bacteria, *Shigella flexneri*. VirF has been recognized as the main transcriptional regulator involved in the virulence of this pathogen. Lack of this VirF mRNA (in the bacteria lacking TGT) reduces the virulence of the bacteria. It has been found that VirF mRNA is a substrate of the *E.coli* TGT, since it has extremely high sequence similarity with *shigella* TGT, and that this recognition causes a site-specific modification of a base in the VirF mRNA and modulates its translation [33]. Based on this information, we cannot rule out that EhTGT and queuine work via a mechanism unrelated to tRNA modification.

To conclude, we have investigated for the first time the role of queuine, a compound produced by the gut microbiota, on the physiology of *E. histolytica*. We showed that queuine is incorporated in the tRNA of the parasite and that it prepares the amoeba to resist oxidative stress *in vitro*. We have also shown that the presence of queuine leads to hypermethylation of tRNA^Asp^_GUC_ in the organism and this agrees with other studies indicating that the role of queuine in methylation is conserved across all species and that its presence may influence Dnmt2 activity. Our study also revealed the role of queuine in shaping the transcriptome of the parasite. Our data also support the role of queuine in the resistance of the parasite to OS but this comes at a cost in terms of virulence. Anti-virulence therapeutic strategies have been proposed to fight against drug-resistant pathogens [85]. In this regard, queuine provided directly to the patient or by selected probiotics may represent a new therapeutic approach for the treatment of amoebiasis.

## Material and methods

### *E. histolytica* culture

*E. histolytica* trophozoites HM-1:IMSS strain were grown under axenic condition at 37 °C in TYI-S-33 medium prepared according to a previously reported protocol [86]. The trophozoites were harvested during the logarithmic phase of growth by chilling the culture tubes at 4 °C and pelleted by centrifugation at 500 g for 5 min. The pellet was washed twice with ice-cold phosphate-buffered saline. Trophozoites were grown with 0.1 μM queuine (a kind gift from Dr. Klaus Reuter, University of Marburg) for 3 days.

### Bacterial strains

The bacterial strains used in this study are *E. coli* BL21(DE3)pLysS and Dh5 alpha. *E. coli* was grown at 37°C in Luria-Bertani (LB) medium [87].

### Cultivation of HeLa cells

HeLa cells (a kind gift from Dr. Kleinberger, Faculty of Medicine, Technion) were maintained in continuous culture using a previously described protocol [88].

### Transfection of *E. histolytica* trophozoites

The transfection of *E. histolytica* trophozoites was done using a previously described protocol [89].

### Effect of queuine on hydrogen peroxide (H_2_O_2_) resistance

*E. histolytica* trophozoites were grown with and without queuine (0.1 μM) for 3 days at 37 °C. Trophozoites were harvested and (1×10^6^) exposed in TYI-S-33 medium (without serum) to various concentrations of H_2_O_2_ (0-5 mM) for 30 min at 37 °C. The viability of the trophozoites was determined with eosin dye exclusion method [78], with this value used to calculate the median lethal dose (LD_50_) of H_2_0_2_.

### Measurement of the cytopathic activity of *E. histolytica*

The cytopathic activity of *E. histolytica* trophozoites was determined using a previously described protocol [90]. Briefly, *E. histolytica* trophozoites (2.5×10^5^ cells/ per well) that were grown with and without queuine, were exposed or not to 2.5mM H_2_O_2_ for 30 min in TYI-S-33 medium (without serum). After the incubation, the cells were resuspended in fresh TYI-S-33 medium (without serum) before their incubation with HeLa cell monolayers in 24-well tissue culture plates at 37°C for 40 min or 90 min. The incubation was stopped by placing the plates on ice and unattached trophozoites were removed by washing the plates with cold Phosphate Buffer Saline (PBS) supplemented with 1% Galactose. The cells were fixed with 4% formaldehyde for 10 min and finally washed twice with cold PBS. HeLa cells remaining attached to the plates were stained with methylene blue (0.1% in 0.1 M borate buffer, pH 8.7). The dye was extracted from the stained cells by 0.1 M HCl, and its color intensity was measured spectrophotometrically at OD_660_. The amount of dye extracted was proportional to the number of intact cells that remained attached to the tissue culture well. The amount of dye extracted from monolayers of tissue-cultured cells that had not interacted with trophozoites served as control (100% of monolayer integrity). The intensity of color was measured in a spectrophotometer (at 660 nm) after appropriate dilution with HCI (0.1 M). Experiments were carried out in duplicate, repeated at least twice, and standard error determined. To decide the exact time at which to terminate an experiment, the progress in tissue culture damage during the incubation with trophozoites was also monitored by periodic observation under an inverted microscope.

### Primers used in this study

The primers used in this study are listed in Table 4.

**Table 4:**
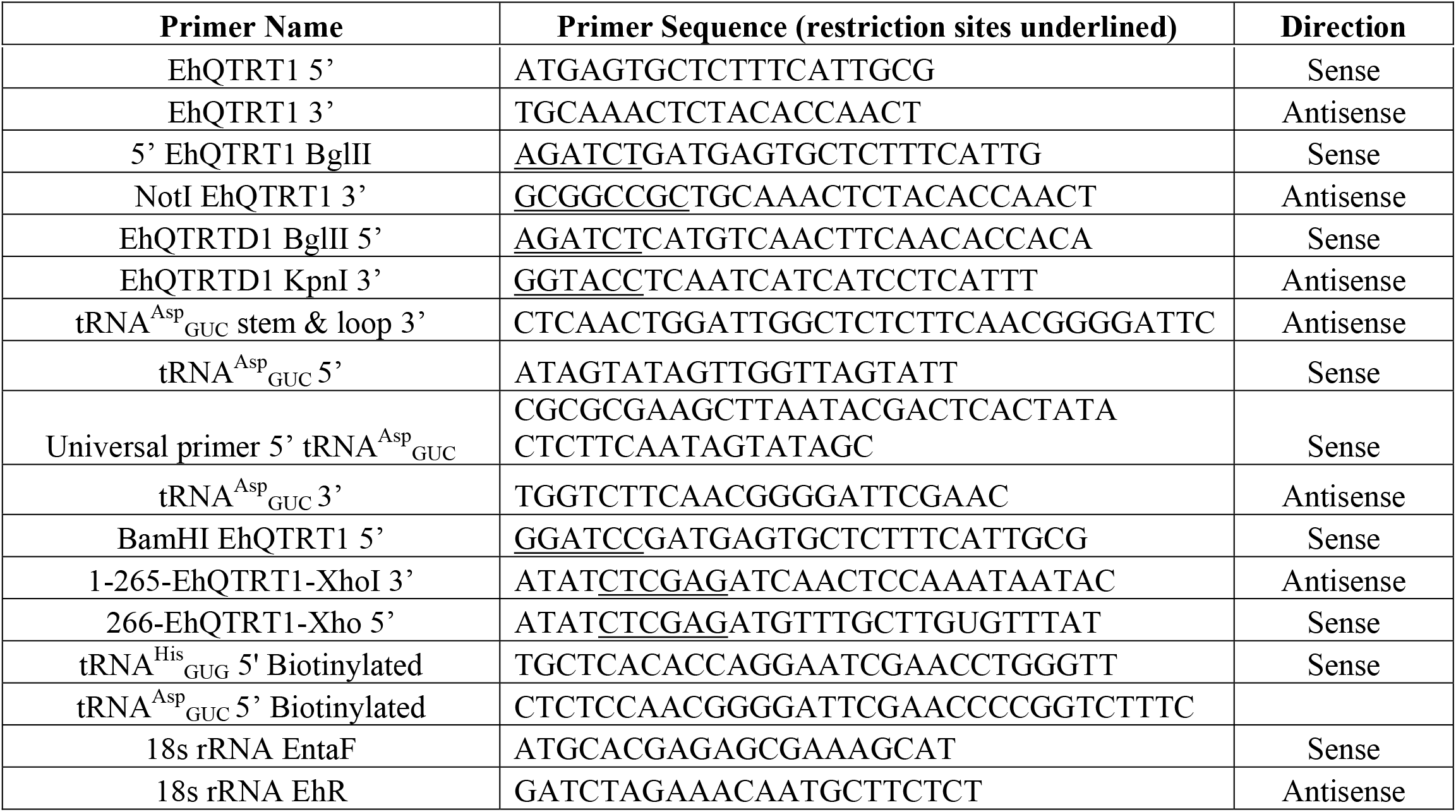
List of all the primers used in this study

### Ethics statement

Animal procedures dealing with the mouse model of intestinal amebiasis were approved by the Institutional Animal Care and Use Committee (No. 211075) and conducted at the AAALAC-accredited National Institute of Infectious Diseases, Japan. Animal procedures dealing with the production of EhTGT antibody were approved by the Technion Animal Care and Use Committee (IL-115-10-16) and conducted at the accredited Faculty of Medicine-Technion, Haifa, Israel.

### Detection and quantification of *E. histolytica* in infected mice stool

The virulent trophozoites were passaged *in vivo* through the CBA/J mouse cecum (purchased from Jackson Laboratory, Japan)[91]. For all intracecal injections, 1 x 10^6^ axenic trophozoites with or without 0.1 μM queuine treatment for three days were inoculated into the proximal and apical region of the cecum [92]. To quantify the number of *E. histolytica* trophozoites in stool, real-time quantitative PCR was performed by using Fast SYBR Green Master Mix (ThermoFisher Scientific, USA) in the StepOne Plus Real-Time PCR System (Applied Biosystems, USA). Primer sets were specific to *E. histolytica* 18S rRNA (EntaF and EhR) [78, 93]. A standard curve was generated using the DNA extracted from trophozoites serially diluted from 10^5^ to 10^0^. Using the standard curve and the stool weight, the number of trophozoites/mg stool was calculated.

### Construction of the pETDuet-1 His-tagged EhQTRT1-EhQTRTD1 vector

For the construction of the pETDuet-1 His-tagged EhQTRT1-EhQTRTD1 vector, EhQTRT1 was amplified from *E. histolytica* genomic DNA with the primers BamHI EhQTRT1 and NotI EhQTRT1. The PCR product was cloned in a pGEM-T easy vector, digested with BamHI and NotI, and then subcloned in a pDuet expression vector that was previously linearized with BamHI and NotI to generate the pETDuet-1 His-tagged EhQTRT1 vector. EhQTRTD1 (EHI_079900) was then amplified from *E. histolytica* cDNA by using primers BglII EhQTRTD1 and KpnI EhQTRTD1 primers, and then cloned in pGEM-T easy vector, digested with BglII and KpnI, and then subcloned in the pETDuet-1 His-tagged EhQTRT1 vector that was previously linearized with BglII and KpnI.

### Construction of the pETDuet-1 his-tagged mEhQTRT1-EhQTRTD1 vector

For the construction of the pETDuet-1 his-tagged mEhQTRT1-EhQTRTD1 having a point mutation of D267 to A267 in EhQTRT1, EhQTRT1 was amplified from *E. histolytica* genomic DNA using primers Standard EhQTRT1 and 1-265-EhQTRT1 XhoI. The second part of EhQTRT1 was amplified using genomic DNA using primers 266-EhQTRT-XhoI (where the Aspartate was replaced with Alanine in the primer) and Reverse Primer EhQTRT1. Both PCR products were purified using QIA PCR purification kit (Qiagen), and digested with XhoI for 1 hour at 37°C and purified. The purified products were ligated at 16°C overnight using T4 ligase. Following ligation, the product was amplified by PCR using primers BamHI EhQTRT1 and NotI EhQTRT1. The PCR product was cloned in pGEM-T easy vector, digested with BamHI and NotI, and further subcloned into pETDuet-1 vector that was previously linearized with BamHI and NotI. Following this, the EhQTRTD1 gene was cloned into pETDuet-1 vector as described above. The resulting plasmid was sequenced to ensure the presence of the desired mutation.

### Construction of silenced EhQTRT1 vector

For the construction of siEhQTRT1 vector, the entire protein-coding region of EhQTRT1 was amplified from *E. histolytica* genomic DNA with 5’ EhQTRT1 BglII and 3’ EhQTRT1 XhoI primers. The resulting PCR product was cloned in pGEM-T easy vector system (Promega), digested with BglII and XhoI, and then subcloned in pEhEx-04-trigger vector containing a 142bp trigger region (EHI_048660) (a kind gift of Dr. Tomoyoshi Nozaki, University of Tokyo, Japan) to generate siEhQTRT1.

Plasmids were sequenced to ensure the presence of unwanted mutations.

### Growth Rate Assays

Trophozoites (3X10^3^/ml) were grown in a 13 ml tube in TYI-S-33 medium at 37 °C and the number of viable trophozoites was counted at 24, 48, and 72 h, respectively by using the eosin dye exclusion method.

### Preparation of recombinant EhTGT complex

Recombinant EhTGT complex was expressed as His-tagged protein in *E. coli* BL21(DE3)pLysS competent cells, which were transfected with the pETDuet-1 vector derived plasmids. The overnight culture was supplemented with Ampicillin (100 μg/ml) and grown at 37 °C. The culture was grown with the addition of 100 μM of ZnSO_4_ until the OD_600_ reached 0.6. The synthesis of the His-tagged protein complex was initiated by adding isopropyl β-D-1-thiogalactopyranoside (IPTG) to the culture at a final concentration of 1 mM. After overnight incubation in the presence of IPTG at 22 °C, the bacteria were harvested and lysed as described elsewhere [94]. The proteins were then eluted with elution buffer (50 mM NaH_2_PO_4_, pH 8.0, 300 mM NaCl, and 250 mM imidazole).

The eluents were further analyzed by SDS-PAGE and the fractions containing EhQTRT1-EhQTRTD1 were combined and then dialysis was performed in dialysis spin columns with a dialysis buffer (25mM HEPES, pH 7.4, 300mM NaCl, 2mM DTT). Two rounds of dialysis were performed for 4 hours each, one at room temperature and the second round of dialysis at 4°C. The proteins were combined, concentrated and then applied to a Superdex 200 increase 10/300 GL column for size-exclusion chromatography. Following their elution from the column with the elution buffer (25mM HEPES, pH 7.3, 100mM NaCl, 50μM ZnSO_4_, 5% glycerol, and 2mM DTT), the eluted fractions (corresponding to the peak) were analyzed by SDS-PAGE, followed by silver staining. The same procedure was used for the purification of mEhQTRT1-QTRTD1.

### Preparation and purification of tRNA^Asp^_GUC_ and tRNA^Val^_CAC_

A commercially synthesized DNA oligomer containing the tRNA^Asp^_GUC_ sequence of *E. histolytica* and the T7 promoter sequence was amplified by using PCR using primers-Universal primer 5’ tRNA^Asp^_GUC_ and tRNA^Asp^_GUC_ 3’ under the following conditions: under the following conditions: primers (20 pmol each), tRNA^Asp^_GUC_ template (500 ng), Mg^2+^ (2 mM), dNTPs (0.5 mM each), ThermolPol Taq buffer (1×), Taq DNA polymerase (2 U), in a final volume of 50 μL. The samples were treated at 94 °C (5 min), followed by 29 cycles in the following order-94 °C (45 s), 60 °C (45 s), 72 °C (45 s). A final extension reaction was allowed to occur at 72 °C for 10 min. The PCR product was purified by using the QIAquick PCR purification kit (Quiagen). The tRNA^Asp^_GUC_ was subsequently generated by in vitro transcription. The conditions for a 50μl reaction are as follows: T7 RNA Pol Buffer (10X)-5μl, rNTPs (25mM each)-4μl, Template-300-400ng, T7 RNA Polymerase-4μl, and purified water up to the final volume. The reaction was incubated overnight at 37°C, after which white precipitates (magnesium pyrophosphate) were removed by centrifugation (13,000 rpm, 5 min). DNAse I was added to remove any unwanted DNA templates in the reaction and incubated for another 30 minutes. The tRNA transcript was ethanol-precipitated at −20°C and then pelleted by centrifugation (20,000x*g*, 30 min, 4 °C). The resulting pellet was resuspended in DEPC water and the concentration was measured using a nanodrop (Thermo Scientific).

tRNA^Val^_CAC_ (EHI_034860) was synthesized as an ultramer desalted RNA oligo (86.2 μg total) by Syntezza Bioscience (Israel).

### Enzymatic assay with [^14^C] guanine

The enzyme activity of the EhTGT complex was performed as described elsewhere with some modifications using 8-[^14^C]-guanine to quantify activity by scintillation counting [39]. Briefly, the kinetic assays were set up with 3μg of *in vitro* transcribed tRNA^Asp^_GUC_ or with 3μg of synthetic tRNA^Val^_CAC_, 0.5μl of 8-[^14^C]-guanine (57mCi/mmol), 0-15μg of EhTGT enzyme, and HEPES reaction buffer (100mM HEPES, pH 7.3, 20mM MgCl_2_, and 5mM DTT) to a final volume of 50μl. Studies were performed in triplicate and samples were incubated at 37°C. Aliquots were taken every 30 minutes for a period of 2 hours and quenched in 3ml of 5% trichloroacetic acid (TCA) on Whatman-glass fiber filters (GE healthcare) and washed 3 times with 1ml ethanol to dry the filter. The filters were then added to a scintillation vial containing 3 ml of scintillation fluid and the samples were analyzed in a scintillation counter where counts were reported in CPM.

### Production of EhTGT antibody

Female BALB/c mice have been injected intraperitoneally with 700 μg recombinant His-tagged TGT protein that was emulsified in complete Freund’s adjuvant (Sigma-Aldrich). Two weeks later, the mice were injected with 700 μg of His-tagged TGT complex protein in incomplete Freund’s adjuvant (Sigma-Aldrich). One week after the 6-week injection, the mice were anesthetized and their blood was collected as described elsewhere [94]. The serum that was obtained from mice that were not injected with His-tagged EhTGT complex was used as the control.

### Western blotting

Total protein extracts of *E. histolytica* trophozoites were prepared according to a published method [95]. Proteins (40 μg) in the total extract were resolved on a 10% SDS-PAGE in SDS-PAGE running buffer (25 mM Tris, 192 mM glycine, 0.1% SDS). The resultant protein bands were visualized after staining with Ponceau red stain. Next, proteins were electrotransferred in protein transfer buffer (25mM Tris, 192mM glycine, 20% methanol, pH 8.3) to nitrocellulose membranes (Whatman, Protran BA83). The blots were first blocked using 3% skim milk and then probed with 1:1000 mouse polyclonal EhTGT (complex) antibody or 1:1000 mouse monoclonal Actin antibody, clone c-4 (MP biotechnologies, CA, USA) for 16 h at 4 °C. After incubation with one of the previously described primary antibodies, the blots were incubated with 1:5000 secondary antibody for one hour at RT (Jackson ImmunoResearch), and then developed using enhanced chemiluminescence (Bio RAD).

### SUnSET assay to measure protein synthesis

Trophozoites (2 × 10^6^/ml) were grown with and without queuine ((0.1μM) for 3 days at 37°C) were exposed or not to 2.5mM H_2_O_2_ for 20 min at 37°C. After treatment with H_2_O_2_, the trophozoites were incubated with 10 μg/ml puromycin (Sigma) for 20 min. For pretreatment of the trophozoites with cycloheximide (Sigma), the trophozoites were incubated with 100 μg/ml cycloheximide for 5 min before the addition of puromycin. The trophozoites were lysed with 1% Igepal (Sigma) in phosphate-buffered saline (PBS) with protease inhibitors. Puromycin was detected by immunoblotting as described above with a 12D10 clone monoclonal puromycin antibody (Millipore). The amount of total protein in each lane was determined by using the No-Stain Protein Labeling Reagent (Thermo Fisher Scientific). Imaging for puromycin and total protein signals was performed on a FUSION FX7 EDGE Imaging System (Witec AG). Quantification of signal density was performed using ImageJ.

### RNA extraction

Total RNA was extracted from control trophozoites, trophozoites treated with queuine (0.1μM for three days at 37°C), and then exposed (or not) to H_2_O_2_ (in three biological replicates) using the TRI reagent kit. according to the manufacturer’s instructions (Sigma-Aldrich). Libraries were built using a Truseq mRNA-Seq Library Preparation Kit (Illumina), according to the manufacturer’s recommendations. Quality control was performed on an Agilent Bioanalyzer. Sequencing was performed on a HiSeq 2500 system (Illumina) and produced 65-base single-end reads.

### RNA-Seq data analysis

Bioinformatics analysis was performed using the RNA-seq pipeline from Sequana [96]. Reads were cleaned of adapter sequences and low-quality sequences using cutadapt version 1.11 [97]. Only sequences at least 25 nt in length were considered for further analysis. Bowtie version 1.2.2 [98], with default parameters, was used for alignment on the reference genome (*Entamoeba histolytica*, from amoebaDB). Genes were counted using featureCounts version 1.4.6-p3 [99] from the Subreads package (parameters: -t gene -g ID -s 1).

Count data were then analyzed using R version 3.5.3^12^ and the Bioconductor package DESeq2 version 1.22.2^29^. Normalization and dispersion estimation were performed with DESeq2 (using the default parameters) and statistical tests for differential expression were performed applying the independent filtering algorithm. A generalized linear model including (i) the effect of stress (+OS vs - OS), (ii) the presence of queuine (Q vs. T) and (iii) the interaction term was set up to test for inter-condition differences in expression and to test the interaction between stress and queuine. For each pairwise comparison, raw p-values were adjusted for multiple testing using the Benjamini and Hochberg procedure [100]. Genes with an adjusted p-value < 0.05 and a fold change > 2 were considered to be differentially expressed.

Principal component analysis and hierarchical clustering were performed using variance-stabilizing transformed counts. Hierarchical clustering of the genes was based on the correlation distance and the Ward aggregation criterion. The average profile was established by computing the mean log2-normalized count over the genes in each of the four identified clusters.

The functional classification, GO term analysis and protein class analysis were performed with PANTHER tools (http://pantherdb.org) [101].

To determine the Gene-Specific Codon Usage (GSCU) for each transcript upregulated in trophozoites cultivated in the presence of queuine, we obtained the protein-coding sequences for 7930 *E. histolytica* genes from the Amoeba database (https://amoebadb.org/amoeba/app/) The *E. histolytica* open reading frames were each analysed using GSCU scripts and methodology previously reported [102]. In brief, GSCU analysis determined the total number of times each codon was used in each gene. GSCU analysis also determined the frequency that each codon was used in each gene, with the codon frequency relative to the other synonymous codons specific to each individual amino acid. For example, the summed frequency for the two glutamic acid codons (GAA and GAG) potentially used in a gene equal 1.0. Note that methionine and tryptophan codons are omitted from GSCU analysis, as they each only have 1 specifying codon and the GSCU frequency is always 1.0 for AUG (methionine) and 1.0 for UGG (tryptophan). Whether a gene was over- or under-represented with a specific codon from a synonymous set, relative to the genome average, was determined by calculating a codon specific Z-score, as described [102]. Gene-specific codon usage data (Z-scores) was analyzed by hierarchical clustering using Cluster 3.0 and visualized as a heat map using Treeview. A spreadsheet containing the gene-specific codon usage data is presented in Table S5.

### Immunofluorescence microscopy

*E. histolytica* trophozoites were transferred to microscope slides as described elsewhere [90]. The attached trophozoites were washed in warm (37°C) PBS, fixed with methanol and then permeabilized with FACS buffer (2% bovine serum albumin in PBS that was supplemented with 0.1% tween) as described by [90]. The slides were then blocked with 5% Goat serum in FACS buffer for 15 min at ambient temperature. The samples were then probed with 1:1000 polyclonal mouse EhTGT (complex) antibody. At the end of the incubation, the slides were washed 3 times in FACS buffer, and then incubated with a 1:250 Alexaflour (Jackson ImmunoResearch) for 3 h at 4 °C. At the end of the incubation, the nuclei of the trophozoites were stained with 1:1000 4’, 6-diamidino-2-phenylindole (DAPI) (Sigma-Aldrich). The samples were then washed with FACS buffer and mounted onto microscope slides with mounting medium (Dako). The specimens were then examined under a confocal immunofluorescence microscope (ZEISS-LSM700 Meta Laser Scanning System confocal imaging system) with a 63X oil immersion objective. For densitometry analysis, the intensity of 15 trophozoites per condition was quantified using image J and signal intensities were normalized to the wild type trophozoites.

### Northern blot analysis

Total RNA (10 μg) were suspended in RNA sample buffer comprising of 17.5μl of RNA mix (10 ml RNA Mix: 600μl of 10X MOPS, 2.1 ml Formaldehyde, 6 ml Formamide, 120 μg/ml ethidium bromide) and 2 μl of RNA loading buffer. Samples were gently mixed and incubated at 70°C for 5 min, then cooled immediately on ice and loaded into 1% agarose gel (6.66% formaldehyde in 1X MOPS buffer). The RNA was blotted on to nylon membrane (Amershan Hybond-N) overnight, the membrane was washed briefly in 1xMOPS and cross-linked by UV using 120 milli joules (Stratagene UV linker). Membranes were hybridized overnight with ^32^-P labeled EhQTRT1 or enolase probes. The membrane was washed with two washing buffers (Washing buffer 1:40mM NaP, 1mM EDTA, 5%SDS Washing buffer 2: 40mM NaP, 1mM EDTA, 1% SDS). Images were developed using a phosphorimager and quantification of signal density from two independent experiments was performed using ImageJ.

### Detection of queuosine in tRNA by chromatography-coupled tandem quadrupole mass spectrometry (LC-MS/MS)

*E. histolytica* trophozoites were grown with and without queuine (0.1μM for three days) and total RNA was isolated using the TRI reagent kit according to the manufacturer’s instructions (Sigma-Aldrich). Three biological replicates from each treatment condition were processed. Samples were processed as blind samples. tRNA was then purified by size-exclusion HPLC performed using a Bio SEC-3 300Å, 7.8 × 300 (internal diameter X length, mm) with 100 mM ammonium acetate pH 7.4 as the mobile phase. The purity of isolated tRNA species was assessed on a Bioanalyzer 2100 system. The average yield of purified tRNA collected ranged from 0.6 to 0.8 mg per sample. Purified tRNA (5μg) from each sample was hydrolyzed in a digestion cocktail containing 10 U benzonase, 5 U bacterial alkaline phosphatases, 0.05 U phosphodiesterase I, 50 μM desferroxamine, 0.5 μg pentostatin, 50 μM butylated hydroxytoluene, 0.25 μg tetrahydrouridine, and 2.5 nmol of the internal standard [^15^N]_5_dA in 10 mM Tris buffer, pH 7.9. The digestion mixture was incubated at 37 °C for 3 h. Digestion mixtures were subsequently removed by passing the mixture through 10 kDa MWCO spin filter. Samples were lyophilized and reconstituted at a final concentration of 100 ng/μl and analyzed by LC-MS/MS. The ribonucleosides were resolved on a Thermo Scientific Hypersil aQ GOLD column (2.1 mm x 100 mm; 1.9 μm particle size) using a mobile phase gradient consisting of 0.1% formic acid as solvent A and acetonitrile with 0.1% formic acid as solvent B at flow rate of 0.3 mL/min and 25 °C using the following elution gradient for solvent B: 0-12 min, 0%; 12-15.3 min, 0-1%; 15.3-18.7 min, 1-6%; 18.7-20 min, 6%; 20-24 min, 6-100%; 24-27.3 74 min, 100%; 27.3-36 min 100-0%; 36-41 min, 0%. The coupled 6490 Triple Quadrupole LC/MS mass spectrometer was operated with the following optimized source parameters: gas temperature, 50 °C; gas flow, 11 L/min; nebulizer pressure, 20 psi; sheath gas temperature, 300 °C; capillary voltage, 1800 V; nozzle voltage, 2000 V; iFunnel RF voltage, 150/50 V and fragmentor voltage, 380 V. Based on a synthetic standard, Q was detected at a retention time of 11.5 min and identified based on two transitions: precursor *m/z* 410, products *m/z* 163 and 295. Quantitation of Q was achieved by normalizing the area under curve (AUC) for the signal intensity of Q against the total AUC for canonical ribonucleosides U, C, A and G for each injected sample. Analysis of each sample condition was performed with 12 biological replicates and data presented as mean and standard error (SEM).

### Bisulfite sequencing of tRNA^Asp^_GUC_

Total RNA isolation and bisulfite conversions were done using a previously described protocol (Schaefer et al. 2009). Bisulfite-treated tRNAs were reverse transcribed by using a tRNA 3’-specific stem-loop primer and amplified with primers that bind only to the deaminated sequences at the 5’ end (Universal primer and a specific tRNAAspGUC stem-loop primer). Amplicons (20) under each condition were subcloned in pGEM-T Easy (Promega) and sequenced (Multi-Disciplinary Laboratories Unit, Bruce Rappaport Faculty of Medicine, Technion).

### APB northern blot to detect tRNA^His^_GUG_

Acryloyl aminophenylboronic acid gels were prepared and run with a few modifications according to Igloi and Kossel [103]. Briefly, 15 μg of RNA was deacetylated in 100 mM Tris-HCl pH 9 for 30 min at 37 °C. RNA was ethanol-precipitated and resuspended in 10μl DEPC water and 1× RNA-loading dye (Fermentas). Samples were then denatured for 10 min at 70 °C and run at 4 °C on Tris Acetate EDTA (TAE), 8 M urea, 15% acrylamide, and 5 mg/ml aminophenylboronic acid (Sigma) on Bio-RAD mini gels. The gel was run at 4°C at 75V for 7 hours until the bromophenol blue reached the bottom of the gel. The gels were then stained with ethidium bromide in 1X TAE for 20 mins and then visualized for equal loading of samples. The gels were destained with ultra-pure water for 20 minutes and were transferred to Hydrobond-XL-membrane (GE healthcare) and electro transferred using 0.5X TAE as the transfer buffer for 45 minutes at 150 V. The membrane was crosslinked using UV using 120 milli joules (Stratagene UV linker) and hybridized twice for 15 minutes each in 5ml hybridization buffer (20mM Sodium phosphate buffer, pH 7.3, 300mM NaCl, 1% SDS), followed by the addition of 150μg/mL heat denatured salmon sperm DNA (ssDNA) to the hybridization buffer and blocked for 1 hour at 60°C. The membrane was then incubated with 15pmol of biotinylated tRNA probes prepared against tRNA^His^_GUG_, and incubated at 60°C for 16 hours. The membrane was then washed for 10 minutes with 5ml wash buffer (20mM Sodium phosphate buffer, pH 7.3, 300mM NaCl, 2mM EDTA, 0.5% SDS) at 60°C, and then incubated in hybridization buffer once at RT for 10 minutes. The membrane was then incubated in streptavidin-HRP conjugate in 5ml hybridization buffer (1:5000) for 30 minutes followed by 2 washes for 10 minutes. The membranes were incubated with enhanced chemiluminescence (Bio-RAD) and then covered in plastic wrap with the RNA side facing upwards and the blots were exposed to X-ray films and developed using a film processor.

### Acid denaturing gel electrophoresis for tRNA^Asp^_GUC_

Acidic gels were prepared and run according to a modified protocol [104]. Briefly, 15 μg of RNA was deacetylated in 100 mM Tris-HCl pH 9 for 30 min at 37 °C. Tubes were briefly centrifuged and 5μl of 2X acidic loading dye (8M Urea, 0.1M Sodium acetate, pH 4.8, 0.05% Bromophenol blue, and 0.05% Xylene cyanol) were added to each sample in the tube. The gel was run at 75V for 4-5 hours at 4°C with acid-TAE running buffer. The gels were blotted and hybridized according to the protocol described above with biotinylated probes for tRNA^Asp^_GUC_.

### Modeling of EhQTRT1 and EhQTRTD1

Swiss Model & Phyre servers were used for template assessment & modelling of the homo dimeric interface of the protein structures. The structure of EhQTRT1 was modeled obtained using combination of homology and ab intio model building approaches. The structure was modeled with high confidence using the high-resolution crystal structure of queuine trna-ribosyltransferase (ec 2.4.2.29)2 (trna-guanine (tm1561) from *thermotoga maritima* (PDB 2ASH). The structure of EhQTRTD1 was modeled using the crystal structure of QTRT2, the non-catalytic subunit of murine tRNA-Guanine transglycosylase (PDB 6FV5).

### Modeling of the EhTGT complex

The homodimers models of both the proteins EhQTRT1 and EhQTRTD1 were energy minimized using AMBER 14 molecular dynamics software (https://ambermd.org/index.php). Using the lowest energy structures, we performed the protein - Protein docking experiment using the cluspro server (https://cluspro.org/login.php). We performed docking checking for the all the possible conformations and defining the active site. The cluspro algorithm considers greedy clustering of ligand positions after obtaining the best poses by rotating the ligand using a grid. We have used the best poses from the top two clusters. The images were created using the pymol software (https://pymol.org/2/).

### Data Availability

RNA-Seq data have been deposited at the Gene Expression Omnibus under the accession number GSE142211.

## Acknowledgment

We thank Mrs. Yuko Umeki at National Institute of Infectious Diseases of Tokyo for her technical assistance, the staff of the Microscopy Imaging Laboratory at the Faculty of Medicine, Technion, Haifa for their help with confocal microscopy resources, and their excellent support in image recording and analysis and Ms Udita Roy and Dr. Daniel Kornitzer, Faculty of Medicine, Technion, Haifa for their help with the purification of the recombinant EhTGT by size-exclusion chromatography.

## Additional Information

Supplementary data are available at DRYAD (https://doi.org/10.5061/dryad.rv15dv465).

